# Rationalizing generation of broad spectrum antibiotics with the addition of a primary amine

**DOI:** 10.1101/2021.08.10.455597

**Authors:** Nandan Haloi, Archit Kumar Vasan, Emily Geddes, Arjun Prasanna, Po-Chao Wen, William W. Metcalf, Paul Hergenrother, Emad Tajkhorshid

## Abstract

Antibiotic resistance of Gram-negative bacteria is largely attributed to the low permeability of their outer membrane (OM). Recently, we disclosed the eNTRy rules, a key lesson of which is that the introduction of a primary amine enhances OM permeation in certain contexts. To understand the molecular basis for this finding, we perform an extensive set of molecular dynamics (MD) simulations and free energy calculations comparing the permeation of aminated and amine-free antibiotic derivatives through the most abundant OM porin of *E. coli*, OmpF. To improve sampling of conformationally flexible drugs in MD simulations, we developed a novel, Monte Carlo and graph theory based algorithm to probe more efficiently the rotational and translational degrees of freedom visited during the permeation of the antibiotic molecule through OmpF. The resulting pathways were then used for free-energy calculations, revealing a lower barrier against the permeation of the aminated compound, substantiating its greater OM permeability. Further analysis revealed that the amine facilitates permeation by enabling the antibiotic to align its dipole to the luminal electric field of the porin and while forming favorable electrostatic interactions with specific, highly-conserved charged residues. The importance of these interactions in permeation was further validated with experimental mutagenesis and whole cell accumulation assays. Overall, this study provides insights on the importance of the primary amine for antibiotic permeation into Gram-negative pathogens that could help the design of future antibiotics. We also offer a new computational approach for calculating free-energy of processes where relevant molecular conformations cannot be efficiently captured.

## Introduction

Gram-negative bacteria are becoming increasingly resistant to available antibiotics, and the development of new antibiotics targeting these bacteria has been slow.^1,2^ A major difficulty is designing antibiotics that can readily permeate the dense outer membrane (OM) of Gramnegative bacteria.^3,4^ Due to the impermeability of the OM, effective antibiotics typically permeate through OM porins, which have evolved to allow diffusion of various substrates necessary for bacterial survival. ^3,4^ Limited knowledge on physicochemical properties that enable permeation of antibiotics through the OM porins, however, has severely hindered the development of new antibiotics for several decades.^5,6^ Without an understanding of these properties to motivate intelligent antibiotic design, even an expansive screening of millions of compounds typically does not provide advanceable lead compounds.^7^ Recently, we were able to identify general determinants for antibiotic permeation by directly measuring the accumulation in *E. coli* for a collection of hundreds of diverse compounds.^8^ These determinants, coined as the eNTRy rules,^9^ were low flexibility, low globularity, and the presence of an ionizable nitrogen, typically a primary amine.

The eNTRy rules provide guidelines for a drug development framework to convert Grampositive only antibiotics to broad spectrum antibiotics through the addition of a primary amine to compounds meeting the flexibility and globularity parameters. ^8^ This approach was used to convert 6-deoxynybomycin (6DNM),^10^ a natural product derivative exclusively active against Gram-positive bacteria, into an antibiotic also effective against Gram-negative pathogens by appending an amino methyl group (6DNM-NH3)^8^ (Fig. 1A). Compared to 6DNM, 6DNM-NH3 shows significantly enhanced accumulation in *E. coli* and a 64-fold increase in antibacterial activity. ^8^ More importantly, 6DNM-NH3 has antibacterial activity against a diverse panel of multidrug resistant Gram-negative pathogens such as *A. baumannii, K. pneumonia* and *E. cloacae*.^8^ This same approach has been used to expand the activity of additional Gram-positive only antibiotics, including Debio-1452 and Ribocil C, into Gram-negative active derivatives, Debio-1452-NH3 and Ribocil C-PA.^11–18^

**Figure 1:**
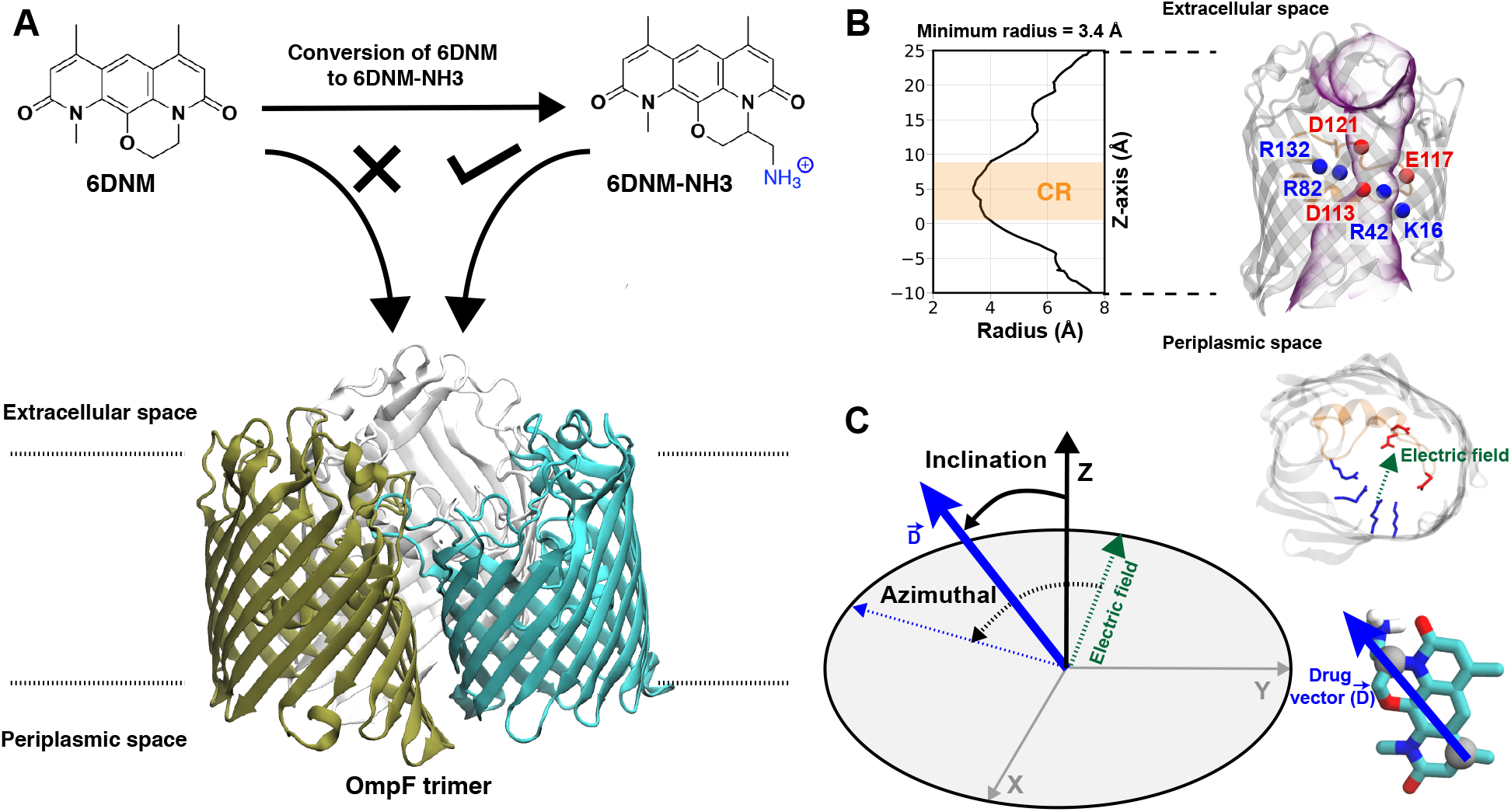
Selective permeability of antibiotics across the *E. coli* porin, OmpF. (A) The conversion of 6DNM to 6DNM-NH3 by introducing a primary amine enhances permeation of the compound through OmpF (each monomer colored differently).^8^ (B, *Top, Left*) The radius profile (calculated using HOLE^39^) of the crystal structure of an OmpF monomer along the membrane normal (*Z*-axis) relative to the membrane center of mass (C.O.M). The constriction region (CR), defined based on a pore radius < 4 Å, is highlighted in orange. The CR spans z-coordinates from 1 to 8 Å. (B, *Top, Right*) Side view of an OmpF monomer, highlighting internal loop L3 in orange and the C*α* atoms of acidic and basic residues within the CR in red and blue, respectively. Purple surface depicts the pore radius profile of OmpF, calculated with the program HOLE.^39^ (B, *Bottom, Right*) Top-down view of OmpF, depicting the the electric field vector (in green dotted arrow), projected onto the membrane plane (*X* – *Y*), generated due to the arrangement of the charged CR residues. The electric field was evaluated using the dipole of all the water molecules within the CR (in 100 ns of equilibrium MD simulation of membrane-embedded OmpF), following a previously described protocol.^25^ (C) Representation of the orientation of 6DNM-NH3 in spherical coordinates. To calculate the orientation angles, a drug vector 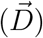 is defined between two carbon atoms (gray spheres), such that the vector points towards the primary amine group. The inclination (*θ*) is the angle between 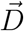 and the membrane normal (*z*-axis); *ϕ* is the angle between the projections of 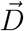 and the electric field vector onto the *XY* plane. The vector for 6DNM is defined using the same two carbon atoms and the angles are calculated using the same manner.

The major proteins responsible for permeating 6DNM-NH3 through the OM of *E. coli* were identified through porin-knockout studies as two trimeric general diffusion *β*-barrel OM porins: outer membrane porin F (OmpF) and OmpC.^8^ In this study, we use OmpF as a model system to investigate how appending a primary amine to 6DNM can significantly enhance its permeation through the OM of *E. coli* using molecular dynamics (MD) simulations and free energy calculations.

The main determinant for the enhanced permeation of 6DNM-NH3 is most likely the favorable interaction of the antibiotic with the porin, especially within the constriction region (CR, Fig. 1B),^19^ a significantly narrowed location within the pore that acts as a bottleneck for antibiotic permeation.^20–31^ Additionally, the CR contains many charged residues including ladders of acidic and basic residues on the L3 loop (a long, internally folded loop) and on the inner barrel wall (Fig. 1B,C), respectively; these residues have been suggested to interact with antibiotics to influence permeation in previous MD simulation and experimental mutagenesis studies.^20–30,32–35^ The arrangement of these charged residues generates a transverse electric field across the CR (Fig. 1B), and alignment of the dipole of antibiotics with this electric field has been suggested to aid in crossing the CR.^24,25,29,36–38^

Despite previous MD simulation studies of antibiotic permeation through OmpF,^20–30,32–35,37,38^ a direct comparison between aminated and amine-free antibiotics and the mechanisms by which the primary amine group aids permeation are lacking. While our previous steered molecular dynamics (SMD) simulations of 6DNM-NH3 translocation through OmpF suggested that addition of the amine group enables interactions with the negatively charged residues of the porin, potentially assisting passage of the antibiotic, ^8^ due to the non-equilibrium nature of SMD and the limited simulation timescale, the findings from these simulations were limited, not allowing for a quantitative description of the process. Therefore, to more comprehensively characterize the molecular mechanisms facilitating permeation of aminated antibiotics, we determined the free-energy landscape associated with translocation of 6DNM and 6DNM-NH3 through OmpF using MD simulations. Calculating these free energies, however, was challenging, because adequately sampling multiple conformations of the drug in the permeation processes using equilibrium or conventional enhanced sampling MD simulations is still computationally intractable. To overcome this challenge, we developed a novel, computationally efficient algorithm, combining Monte Carlo and graph theory, to more efficiently probe the high dimensional permeation pathways of the two antibiotics through OmpF. The resulting pathways were then used for one dimensional bias exchange umbrella sampling (1D-BEUS) simulations.

From 1D-BEUS simulations, we evaluated the free-energy of permeation for 6DNM-NH3 and 6DNM which reveals a lower energetic barrier and greater permeability for the aminated antibiotic through OmpF, in accord with our previous experiments. ^8^ Further analysis revealed that the amine facilitates permeation by enabling 6DNM-NH3 to align its dipole to the luminal electric field of the porin and by forming favorable electrostatic interactions with charged residues while hopping along the pore. The importance of these key interactions in permeation of 6DNM-NH3 was further validated by experimental mutagenesis and whole-cell accumulation assays.

## Methods

### Computational Methods

Here we first provide an overview of our computational approach to obtain free energy landscapes of antibiotic permeation. In principle, Free energies can be evaluated from equilibrium MD trajectory data by measuring the probability of the antibiotic along the pore (*Z*) axis. However, drug permeation occurs on timescales^40^ that may not be sampled sufficiently using equilibrium MD simulations. Therefore, it is necessary to adopt a more rigorous enhanced sampling method such as bias-exchange umbrella sampling (BEUS),^41–45^ metadynamics,^46^ or adaptive biasing force. ^47^ Theoretically, these methods can enhance sampling of all relevant slow degrees of freedoms (DOFs), i.e., those DOFs that cannot be sufficiently sampled by equilibrium MD. However, in practice, multi-dimensional enhanced sampling simulations are computationally prohibitive, and reaching convergence is challenging. ^34,35,45,48–50^ Typically, in an enhanced-sampling simulation, a bias is applied only along the most relevant slow DOF, with the assumption that other orthogonal slow DOFs will be properly sampled during the simulation. A break in this assumption is especially apparent when studying antibiotic permeation through porins. Enhanced sampling methods typically only bias the translational coordinate of the antibiotic along the pore axis (*Z*);^34,35^ however, this does not allow sufficient sampling of other orthogonal slow DOFs, e.g., the drug orientation, since the narrow CR of these porins restricts reorientation. Insufficient sampling of other slow DOFs would provide an incomplete picture of the drug permeation process. Our approach to account for multiple slow DOFs is to use a heuristic algorithm to exhaustively sample the high dimensional conformational space and then use this sampling to determine an optimal initial pathway. This pathway will then serve to generate seeds for one dimensional BEUS (1D-BEUS) simulations. As 1D-BEUS does not exhaustively sample orthogonal slow DOFs, having a proper initial pathway will ensure that the energetics we derive are physically meaningful. A simple approach to generate such an optimal initial pathway is to perform multiple independent steered MD (SMD) simulations and use the lowest nonequilibrium work run. ^44,45^ However, unless a very large number of SMD runs are performed, conformational space will be still insufficiently sampled and the resulting pathway will be based on an incomplete data set.

To address this issue we developed a step-wise approach: (1) create a dataset of discrete poses of the antibiotic by exhaustively exploring the translational and rotational DOFs within the porin using a grid-based workflow; ^51^ (2) use this dataset to determine multiple energetically favorable permeation trajectories from extracellular to periplasmic space using our novel Monte Carlo based pathway search (MCPS) algorithm; (3) determine the most likely permeation pathway sampled in our MCPS trajectories using Dijkstra’s algorithm; and (4) use this most likely pathway to determine energetics of the permeation process using BEUS. ^41–45^ Details of these steps are discussed in the following sections.

### Creation of a dataset with discrete antibiotic poses

A workflow (Fig. 2A), similar to the one developed by us previously, ^51^ was designed to exhaustively explore the translational and rotational DOFs of 6DNM and 6DNM-NH3, independently, within the pore of the crystal structure of OmpF (PDB ID: 3POX).^52^ To improve the computational efficiency of our approach, each pose was generated in vacuum and with monomeric OmpF; however, once the optimal initial pathway is obtained, we refine it using 1D-BEUS in a more realistic, fully solvated membrane-embedded protein-drug system with trimeric OmpF (see below). We will describe this workflow using 6DNM-NH3 as an example; the same procedure was used for 6DNM.

**Figure 2:**
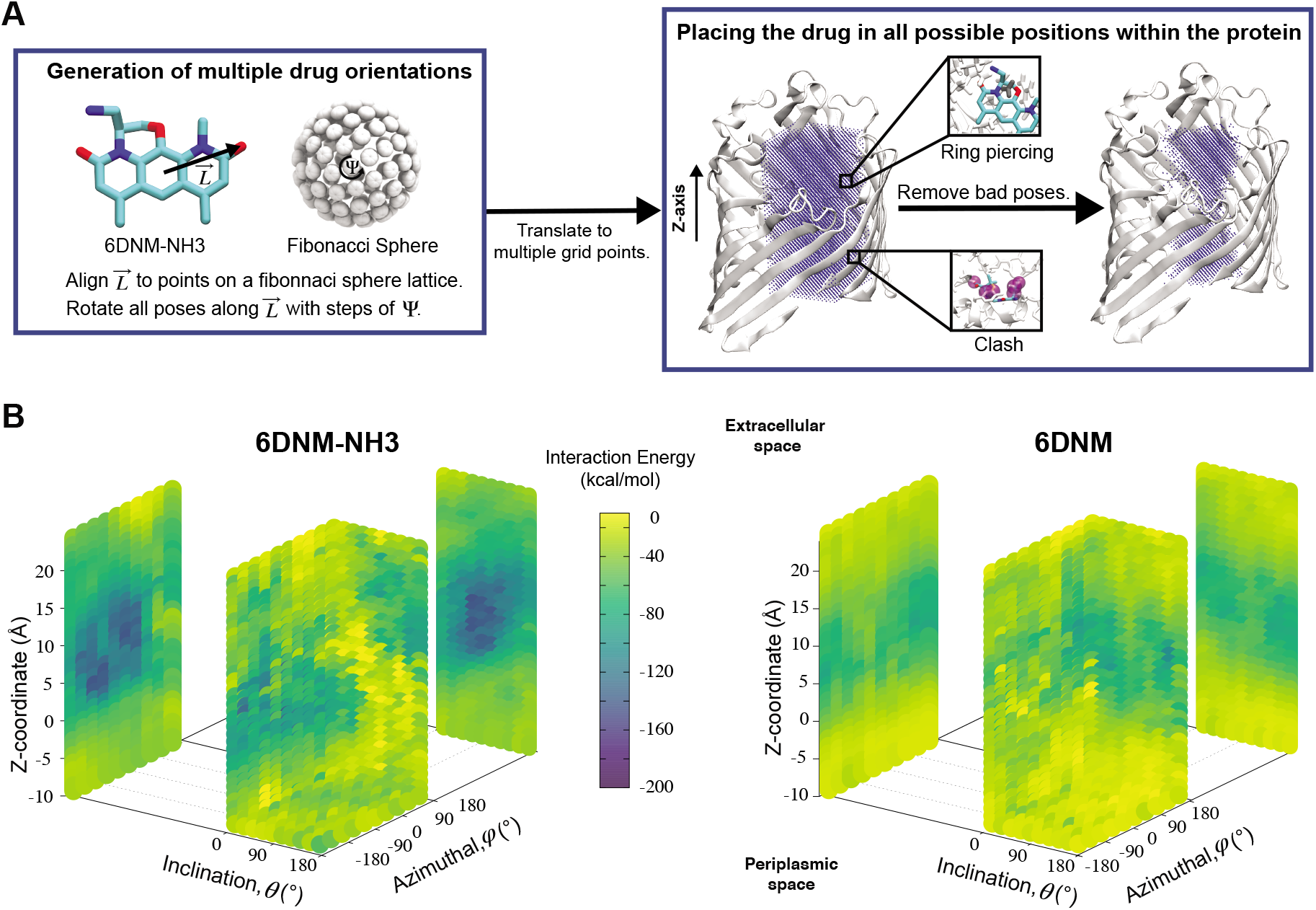
A workflow to create a dataset of discrete antibiotic poses by exhaustively exploring the translational and rotational DOFs within the pore of an OM porin and to evaluate the antibiotic-protein interaction energy (IE) for each pose. (A, *Left*) A set of poses of 6DNM-NH3 with multiple antibiotic orientations was generated by mapping the antibiotic along its longest axis from its C.O.M. 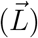 onto a Fibonacci spherical lattice (FSL).^55^ Each of the generated antibiotic poses was further rotated along 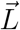, with an interval of Ψ. (A, *Right*) The set of 6DNM-NH3 orientations were placed at every point of a 3-dimensional rectangular grid. During this placement, poses with clashes between antibiotic and the protein or ring piercings were removed. The same process was applied to generate 6DNM poses within OmpF. (B) IE between the protein and poses of 6DNM-NH3 (*left*) or 6DNM (*right*), projected along the *Z*-coordinate, inclination (*θ*) and azimuthal (*ϕ*) of the antibiotic.

First, the residues E296, D312, and D127 in the OmpF crystal structure were protonated. ^36,53,54^ Then, using the center of the CR residues (Lys16, Tyr40, Arg42, Arg82, Arg132, Tyr102, Tyr106, Asp113, Met114, Leu115, Pro116, Glu117, Phe118, Gly119 and Gly120) as the origin, a 16 × 16 × 34 search grid (1 Å spacing) was defined (Fig. 2A). Copies of 6DNM-NH3 were first placed at every grid point, and then each copy was rotated by mapping a vector, 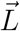 (longest axis from the C.O.M of 6DNM-NH3 to the furthest atom, the oxygen of carbonyl2 [Fig. 4B]) onto a Fibonacci spherical lattice (FSL)^55^ of 25 points, generating multiple orientations at each grid point. During rotation, the C.O.M of 6DNM-NH3 was set to be at the center of the FSL, while the C.O.M of the oxygen of carbonyl2 was arranged on the FSL. Each of the 25 generated orientations at each grid point was then self-rotated along 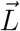, with an interval of Ψ = 45°, creating a total of 200 poses at each grid point. We reason that this approach would evenly survey possible orientations of 6DNM-NH3. This strategy resulted in a total of 1.7 million distinct drug-protein poses.

During our grid-based orientation search process, bad contacts between protein/drug occurred in several poses. These bad contacts were either drug-protein clashes or piercings of the protein into the rings of 6DNM-NH3 (Fig. 2A). Clashes were defined if within a pose, 6DNM-NH3 had a contact distance of <1.0 Å with at least 4 heavy atoms of the protein. Ring piercings were defined by first decomposing rings into triangles and nearby bonds from protein into line segments, and then applying a geometric test to determine if line segments intersected any triangles. Poses associated with clashes and ring piercings were removed, resulting in a total of about 700,000 distinct poses.

Each pose was subjected to 200 steps of energy minimization to relax it to an energetically favorable state. Minimization was performed using the Generalized Born implicit solvent (GBIS) module implemented in NAMD2^56,57^ to account for the effect of solvation on interactions between the drug and protein without including explicit solvent. GBIS calculates molecular electrostatics in solvent by representing water as a dielectric continuum as described by the Poisson Boltzmann equation. The elimination of explicit solvent greatly accelerates modeling efficiency. During minimization, the protein side chains and the drug were allowed to move, while protein backbone was kept fixed in order to allow the drug to relax in its environment without causing a significant conformational change in the protein.

### Monte Carlo based pathway search (MCPS) algorithm

We used the minimized poses (*S_i_*) obtained from the previous step to probe potential permeation pathways from the extracellular to periplasmic space using Monte Carlo (MC). To use MC, we first generated an energy landscape by calculating the drug-protein interaction energy (IE) for each pose (*E*(*S_i_*)) by evaluating the sum of van der Waals (vdW) and electrostatic interaction energies using NAMD2^56,57^ (Fig. 2B). We then walk through this approximate energy landscape along the membrane normal (*Z*) to determine favorable pathways connecting extracellular and periplasmic spaces while also exploring orientation space.

To do this, we first divide the *Z*-coordinate space from the extracellular to the periplasmic space into *N* 1-Å bins, 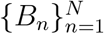 (Fig. 3A,B). Then, we assign poses to these bins, according to the *Z*-coordinate of the C.O.M. of the drug in each pose. That is, a pose, *S_i_*, is assigned to a bin, *B_n_*, if

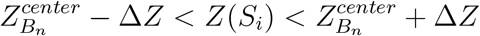

where 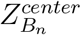, Δ*Z* and *Z*(*S_i_*) refer to the center of *B_n_*, the half-width of *B_n_* (=0.5 Å) from 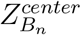, and the *Z*-coordinate of the center of mass (C.O.M.) of the antibiotic at a particular pose, respectively. Then, a random pose (*S*_1_) from a bin closest to the extracellular space, *B*_1_, is selected as the starting point of the trajectory. Now, we need to identify a new pose in the next bin to serve as the next step in the trajectory towards the periplasmic space (decreasing *Z* by 1 Å). Since we are interested in modeling a chemically relevant permeation trajectory, we also need to ensure that the orientation of the antibiotic, defined by the inclination (*θ*) and azimuthal (*ϕ*) angles, does not change substantially in the next step (mechanistically unlikely), since these angular spaces are slow DOFs. To calculate the orientation angles, a drug vector 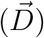 is defined (Fig. 1C), such that the vector points towards the positively charged primary amine group. Then *θ* is calculated as the angle between 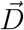 and the membrane normal (*Z*-axis), while *ϕ* is the angle between the projections of 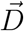 and the electric field vector of the protein onto the *XY* plane (Fig. 1C).

**Figure 3:**
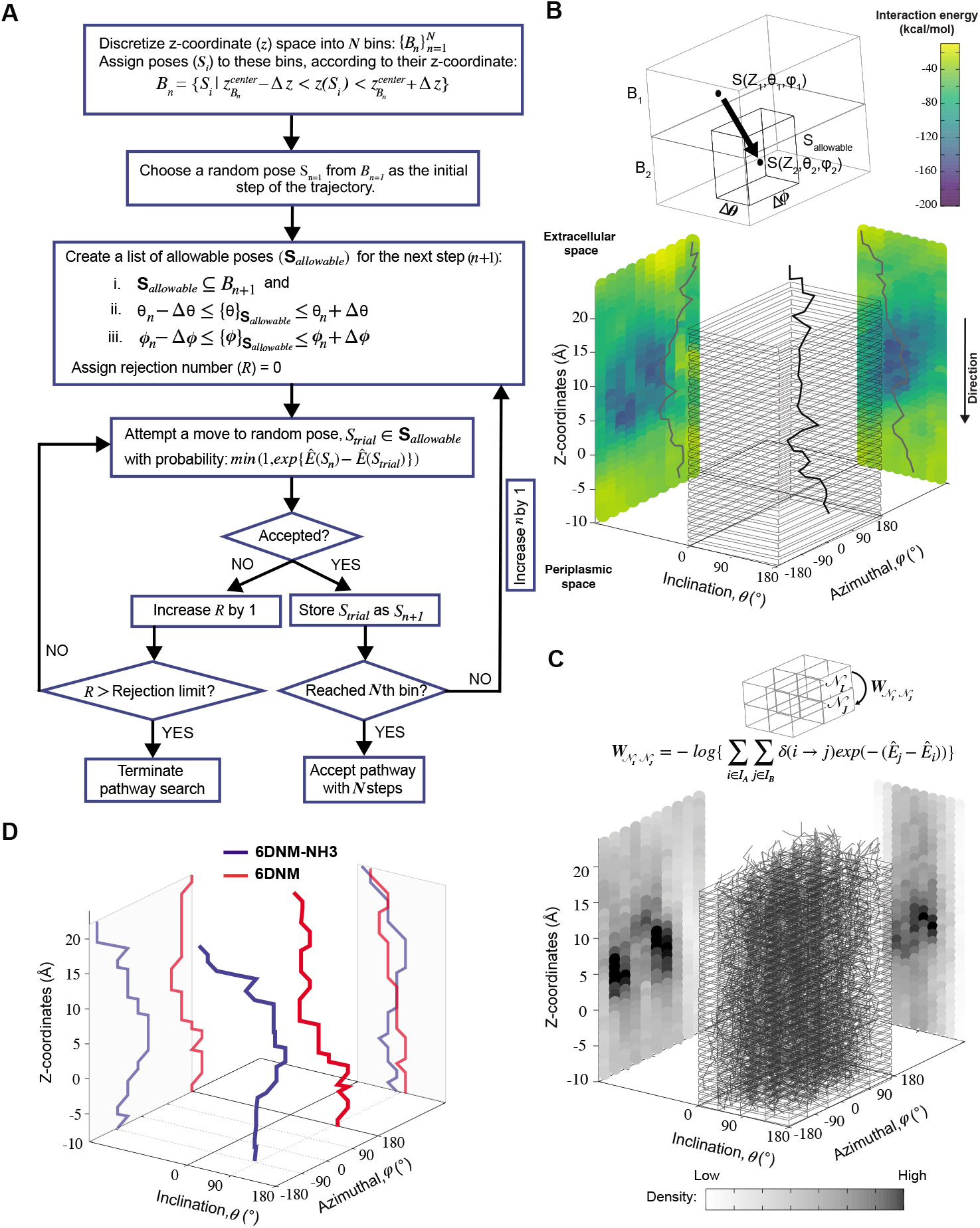
Determination of most likely antibiotic permeation pathways through OM porins using the dataset whose generation is described in Fig. 2. (A) Flowchart describing the Monte Carlo based pathway search (MCPS) algorithm. (B) A representative MCPS trajectory of 6DNM-NH3. To run MCPS, briefly, we first discretize the *Z*-coordinate space into *N* bins 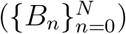 to which we assign the poses generated in Fig. 2A. Then, Monte Carlo (MC) moves are used to walk through the IE landscape generated in Fig. 2B to determine favorable trajectories connecting extracellular and periplasmic spaces. In each MC move, limited change in antibiotic orientation and position are allowed. (C) Generation of a connected graph using multiple MCPS trajectories. Boltzmann weighted densities of the 2000 trajectories are projected along the *Z*-coordinate, inclination (*θ*) and azimuthal (*ϕ*) of the antibiotic. (D) The most likely permeation pathway for 6DNM (red) and 6DNM-NH3 (blue), evaluated using Dijkstra’s algorithm. ^58^

To transition to the next step, we compile an allowable list of poses in the next bin (*B*_2_) with *θ* and *ϕ* within a range (Δ*θ* = 18° and Δ*ϕ* = 36°, respectively) of *θ*(*S*_0_) and *ϕ*(*S*_0_). Within the compiled list, a trial pose, *S_trial_* is chosen randomly and a move is attempted. The move is accepted with probability:

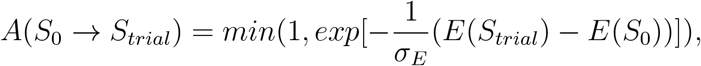

where *σ_E_*, the standard deviation of the IE of all poses, is used to standardize the energies. This standardization ensures that the IE of each pose is weighted according to the energy scaling of the IE distribution since the energy landscape of the system can be of an arbitrary scale. If the move is rejected, then we attempt a new move to another *S_trial_* from the compiled list. Once a move is accepted, *S_trial_* is stored as *S*_1_, the next step in the trajectory, and the protocol is repeated until the last bin, *B_N_*, at the periplasmic space is reached. Then, the series of accepted poses: *S*_1_, *S*_2_, *S*_3_, …, *S_N_* is stored as a permeation trajectory.

### Identifying the most likely pathway

We generated 2,000 MCPS trajectories to exhaustively search for the possible pathways the antibiotic can take to permeate through OmpF. These trajectories resulted in exhaustive and convergent sampling of the conformational space (Figs. S7 and S8). Then, to determine the most likely pathway (that will be used to generate seeds for BEUS simulations) from these trajectories, we build a directed graph and apply Dijkstra’s algorithm. ^58^ In the directed graph, each node 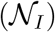 is defined as a set of poses such that:

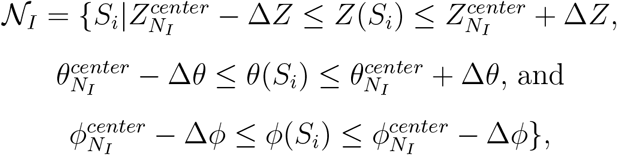

where 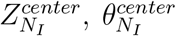, and 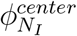 refer to the center of *N_I_* in the *Z*, *θ*, and *ϕ* coordinates, respectively, and Δ*Z*, Δ*θ*, and Δ*ϕ* represent the range from their respective centers (Fig. 3C). Once nodes are defined, edge weights between any two nodes 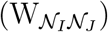 are determined by calculating the negative logarithm of transition counts (each transition weighted by the Boltzmann factor of the two connected poses making up the transition):

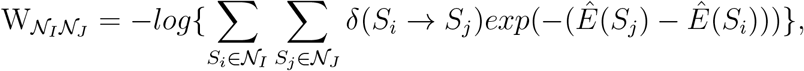

where 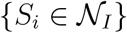 is the set of all poses within node 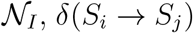 is 1 if a transition occurs between *S_i_* and *S_j_* (otherwise it is 0) and 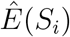 is the standardized energy of pose *S_i_*. The negative logarithm was taken because Dijkstra’s algorithm determines the pathway with the lowest total edge weight from source to sink node. Sink and source nodes are defined as two virtual nodes, *V*_1_ (source) and *V*_2_ (sink), each connected to the set of nodes with maximum (extracellular set) and minimum (periplasmic set) 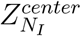, respectively. Each connection was initially assigned an arbitrary edge weight. We then used Dijkstra’s algorithm to compute the pathway connecting *V*_1_ and *V*_2_ with the lowest overall edge weight, resulting in the most likely pathway from extracellular to periplasmic space with *N* nodes: 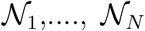, whose centers are separated by 1 Å along the *Z*-coordinate.

### Free-energy calculations using MCPS derived pathways

To investigate the energetics of permeation of 6DNM-NH3 and 6DNM, the lowest energy poses in the nodes along the most likely pathways for each drug were used to serve as windows in 2 independent 1D-BEUS simulations.^41–45^ The most likely path contains poses with *Z*-values ranging from −10 to 24 Å (relative to the midplane of the membrane) of 1 Å width; to obtain reference free energies, we extended the number of windows in the extracellular and periplasmic spaces such that the terminal windows are at least 10 Å away from any atom of the protein. To ensure adequate histogram overlap in BEUS, additional windows were added in between the original windows starting from the entrance to the exit of the CR (*Z* =-3 to 12 Å) such that the window width was 0.5 Å within this region. In total, 94 windows were used spanning 80 Å from the extracellular (*Z*=46 Å) to the periplasmic (*Z* = −34 Å) space.

To take into account the biologically relevant configuration of the system, for each window we built a trimeric, membrane-embedded protein-drug system. To do this, we first aligned the backbone atoms of the *β*-barrel residues of the monomeric protein of each window to those of a single monomer of the X-ray structure of trimeric OmpF (PDB ID: 3POX). ^52^ Then, we merged the resulting coordinates of the aligned antibiotic-monomer system with the two additional monomers of the trimeric OmpF. In the generated trimer for each window, residues E296, D312, and D127 were protonated.^36,53,54^ The windows were then embedded in a symmetric membrane composed of 1,2-dimyristoyl-sn-glycero-3-phosphocholine (DMPC) lipid molecules in each leaflet generated using the Membrane Builder module of CHARMM-GUI. ^59^ We did not use an OM composition containing lipopolysaccharides (LPS) since the membrane composition is unlikely to influence the dynamics of the drug inside the pore. Each window was solvated with TIP3P water^60^ and buffered in 0.15 M NaCl to generate systems containing ~140,000 atoms with dimensions of 120 × 120 × 100 Å^3^.

Before performing 1D-BEUS simulations, each drug-bound, membrane-embedded trimer was minimized using 10,000 steps of the steepest descent algorithm, and then the molecular system was allowed to relax at the center of each window during a 1-ns MD simulation while the drug and heavy atoms of the protein were restrained with a force constant of 1 kcal mol^−1^ Å^−2^. This was followed by 35 ns of 1D-BEUS simulations (until the convergence of the free-energy, Fig. S10) using the distance along the membrane normal (*Z*-axis) between the drug’s C.O.M and the C.O.M of the antibiotic-containing monomer as the collective variable. The force constant of each window was chosen such that the exchange ratio for each window was between 0.15 and 0.3.^61^ The resultant force constants were 2.0 kcal mol^−1^ Å^−2^ for all windows except for windows from the entrance to exit of the CR, which had force constants of 7.0 kcal mol^−1^ Å^−2^. As shown in Fig. S9, using these force constants resulted in good window overlap for each drug. The first 20 ns of each window were discarded as equilibration, and the rest of the trajectories were used in evaluating the free-energy. A non-parametric variation of the weighted histogram analysis method (WHAM), ^62^ proposed by Bartels^63^ and implemented by Moradi and Tajkhorshid^45^ was used to estimate the free-energy profile from the BEUS simulations.

### Steered molecular dynamics (SMD)

To compare the free energies for 6DNM and 6DNM-NH3 along MCPS-derived pathways to those along pathways derived using steered molecular dynamics (SMD), SMD simulations were performed and their trajectories were used to generate windows for two additional 1D-BEUS simulations for each drug. To perform SMD, drug-protein systems were built by first embedding the X-ray structure of trimeric OmpF (PDB ID: 3POX)^52^ in a symmetric membrane composed of DMPC lipids generated using the Membrane Builder module of CHARMM-GUI.^59^ In each monomer, residues E296, D312, and D127 were protonated.^36,53,54^ Each system was solvated with TIP3P water^60^ and neutralized in 0.15 M NaCl to generate systems each containing ~140,000 atoms with dimensions of 120 × 120 × 100 Å^3^. Ten replicas of each drug were then placed at the pore mouth with different initial orientations, resulting in a total of 20 independent drug-protein systems.

The systems were then minimized using the steepest descent algorithm for 10,000 steps, followed by an initial equilibration of 1 ns, during which the heavy atoms of the protein and the drug were harmonically restrained using a force constant of 1.0kcalmol^−1^ Å^−2^. Then, a harmonic spring with a force constant of 10 kcal mol^−1^ Å^−2^ was attached to the C.O.M. of each drug, to pull it along the *Z*-coordinate from the pore mouth on the extracellular side to periplasmic space through a monomer of OmpF at a constant velocity of 0.5 Å ns^−1^. To avoid structural deformation of the protein due to the strong non-equilibrium nature of SMD, protein heavy atoms were harmonically restrained using a force constant of 1.0kcalmol^−1^ Å^−2^.

The SMD trajectory for each drug associated with the lowest non-equilibrium work was then used to generate 1-Å windows, except for the region between the extracellular entrance and the exit of the CR (*Z* =-3 to 12), where 0.5-Å windows were used to ensure adequate histogram overlap. To obtain reference free energies, additional windows were added by extending the lowest-work SMD trajectory of each drug towards the water in the extracellular and periplasmic spaces such that terminal windows were at least 10 Å away from any atom of the protein. In total, 94 windows were generated along the *Z*-coordinate for each drug-protein system. Then, two independent 1D-BEUS simulations were performed using the same protocol described above. This protocol resulted in a good exchange ratio (between 0.15 and 0.3) for each window as suggested previously^61^ and good window overlap (Fig. S2). Free energy of each drug was then evaluated using the non-parametric WHAM. ^45,62,63^

### MD simulation protocol

MD simulations in this study were performed using NAMD2^56,57^ utilizing CHARMM36m^64^ and CHARMM36^65^ force field parameters for proteins and lipids, respectively. The force field parameters for both drugs were generated using the CHARMM General Force Field (CGenFF)^66–68^ with the ParamChem server. Bonded and short-range nonbonded interactions were calculated every 2 fs, and periodic boundary conditions were employed in all three dimensions. The particle mesh Ewald (PME) method^69^ was used to calculate long-range electrostatic interactions every 4 fs with a grid density of 1 Å^−3^. A force-based smoothing function was employed for pairwise nonbonded interactions at a distance of 10 Å with a cutoff of 12 Å. Pairs of atoms whose interactions were evaluated were searched and updated every 20 fs. A cutoff (13.5 Å) slightly longer than the nonbonded cutoff was applied to search for the interacting atom pairs. Constant pressure was maintained at a target of 1 atm using the Nosé-Hoover Langevin piston method.^70,71^ Langevin dynamics maintained a constant temperature of 310 K with a damping coefficient, *γ*, of 0.5 ps^−1^ applied to all atoms. Simulation trajectories were collected every 10 ps.

### Antibiotic permeability calculation

The antibiotic permeability (*P*) was calculated by treating the antibiotic translocation process as a one-dimensional diffusion-drift problem: ^24,72,73^

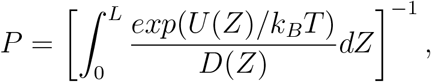

where *U*(*Z*) is the free-energy (calculated as described above) and *D*(*Z*) is the diffusion coefficient along the pore axis (*Z*); *L* is the length of the pore and *k_B_T* is the thermodynamic temperature.

To calculate *D*(*Z*), a set of independent simulations were initiated from the same seeds used for BEUS simulations, during which the *Z* position of the antibiotic was restrained by means of a harmonic potential with a force constant of 10 kcal mol^−1^ Å^−2^ (as suggested by Lee *et al*.^73^). Each simulation was performed for 10ns. From these simulations, *D*(*Z*) was calculated in the framework of an overdamped harmonic oscillator, ^73–75^

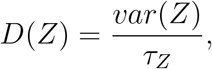

where the variance, *var*(*Z*), is equal to 〈*Z*^2^〉 – 〈*Z*〉^2^, and *τ_Z_* is the characteristic time of the normalized autocorrelation function of *Z*:

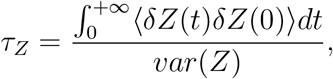

where, *δZ*(*t*) = *Z*(*t*) – 〈*Z*〉.

### Analysis

System visualization and analyses were carried out using VMD. ^76^ Drug-protein interaction energy (vdW + electrostatic) was calculated using the NAMDEnergy plugin of VMD. A hydrogen bond was counted between an electronegative atom with a hydrogen atom (H) covalently bound to it (the donor, D), and another electronegative atom (the acceptor, A), provided that the distance D-A was less than 3Å and the angle D-H-A was more than 120°.

## Experimental Methods

### Mutant construction

*E. coli ompF* mutants were constructed as described before.^77^ Briefly, plasmids carrying the desired mutation were used to replace the wild-type (WT) *ompF* allele in *E. coli* BW25113.^78^ To do this, the mutagenic plasmids were moved from *E. coli* WM6026^79^ donors to BW25113 recipients via conjugation with selection for exconjugants on LB agar with kanamycin (50 *μ*g/ml). Because the plasmids are incapable of replication in *E. coli* BW25113, these exconjugants arise by integration into the recipient chromosome via homologous recombination, creating partial diploids of the *ompF* region, with one WT and one mutant allele separated by the plasmid vector. Kanamycin-sensitive recombinants that had resolved the partial diploid state were then selected by growth on a modified LB medium that lacked NaCl and that contained 5% sucrose. The presence of the desired mutation was verified by DNA sequencing of *ompF* PCR products. *E. coli* WM8633 (*lacI^q^, rrnB3*, Δ*lac4787, hsdR514*, Δ*(araBAD)567*, Δ*(rhaBAD)568, rph-1, ompF(D113F)*) was constructed via allele exchange using plasmid pAP002. *E. coli* WM8636 (*lacI^q^, rrnB3*, Δ*lac4787, hsdR514*, Δ*(araBAD)567*, Δ*(rhaBAD)568, rph-1, ompF(D113V)*) was constructed via allele exchange using plasmid pAP003. The mutagenic plasmids carrying the *ompF* mutations are derivatives of pHC001A.^80^ These plasmids were constructed by HiFi DNA assembly (New England Biolabs, Ipswich, MA) using NotI-digested pHC001A and a mutagenized *ompF* allele produced by overlap-extension PCR. Plasmid pAP002 was constructed using the following primers: ompF-F: **CCGGGGGATCCACTAGTTCTAGAGCGGCCGC**ACGTAACT-GGCGTGCAAAAC; ompF-R: **AAAGCTGGAGCTCCACCGCGGTGGCGGCCGC**AGCGGCGGTAATGTTCTCAA; ompF(D113V)F: TTCGCGGGTCTTAAATACGCT**GTT**-GTTGGTTCTTTCGATTACGGCCGTAA; and ompF(D113V)R: TTACGGCCGTAATCG-AAAGAACCAAC**AAC**AGCGTATTTAAGACCCGCGAA. Plasmid pAP003 was constructed using ompF-F; ompF-R; ompF(D113F)F: TTCGCGGGTCTTAAATACGCT**TTC**GTTGGTT-CTTTCGATTACGGCCGTAA; and ompF(D113F)R: TTTACGGCCGTAATCGAAAGAAC-CAAC**GAA**AGCGTATTTAAGACCCGCGAA. Bold sequences in primers ompF-F and ompF-R represent homology to pHC001A required for HiFi assembly, whereas those in ompF(D113V)F, ompF(D113V)R, ompF(D113F)F, and ompF(D113F)R represent the codon changed to produce the desired mutation.

### Accumulation assay

The accumulation assay^8,81^ was performed in triplicate, with tetracycline as a positive control. The strain *E. coli* BW25113 (WT) and two strains containing single mutants in ompF, *E. coli* ompF^D113V^ and *E. coli* ompF^D113F^, were used for these experiments. For each replicate, 2.5 ml of an overnight culture of *E. coli* was diluted into 250 ml of fresh Luria Bertani broth (Lennox). Cultures were grown at 37°C with shaking to an optical density at a wavelength of 600 nm (OD_600_) of 0.55-0.60. The bacteria were pelleted at 3,220 RCF for 10 min at 4°C, and the supernatant was discarded. The pellets were resuspended in 40 ml phosphate buffered saline (PBS) and pelleted as before, and the supernatant was discarded. The pellets were resuspended in 8.8 ml fresh PBS and aliquoted into 1.7 ml Eppendorf tubes (875 *μ*l each). The number of colony-forming units (CFUs) was determined by a calibration curve. The samples were equilibrated at 37°C with shaking for 5 min, then compound was added (final concentration=50 *μ*M) and samples were incubated at 37°C with shaking for 10 min. A 10-min time point was chosen because it is longer than the predicted amount of time required to reach a steady-state concentration, ^82^ but short enough to minimize metabolic and growth changes (no changes in OD_600_ or CFU count observed). After incubation, 800 *μ*l of the cultures was carefully layered on 700 *μ*l of silicone oil (9:1 AR20 (Acros; catalogue number: 174665000)/ Sigma High Temperature (Sigma–Aldrich; catalogue number: 175633), cooled to −78°C). Bacteria were pelleted through the oil by centrifuging at 13,000RCF for 2 min at room temperature (with the supernatant remaining above the oil). The supernatant and oil were then removed by pipetting. To lyse the samples, each pellet was resuspended in 200 *μ*l water, and then subjected to three freeze–thaw cycle of 3 min in liquid nitrogen followed by 3 min in a water bath at 65°C. The lysates were pelleted at 13,000 RCF for 2 min at room temperature and the supernatant was collected (180 *μ*l). The debris were resuspended in 100 μl methanol and pelleted as before. The supernatants were removed and combined with the previous supernatants collected. Finally, the remaining debris were removed by centrifuging at 20,000 RCF for 10 min at room temperature.

Supernatants were analyzed with the QTRAP 5500 LC/MS/MS system (Sciex) in the Metabolomics Laboratory of the Roy J. Carver Biotechnology Center, University of Illinois at Urbana-Champaign. Software Analyst 1.6.2 was used for data acquisition and analysis. The 1200 Series HPLC System (Agilent Technologies) used included a degasser, an autosampler and a binary pump. The liquid chromatography separation was performed on an Agilent Zorbax SB-Aq column (4.6mm × 50mm; 5 *μ*m) with mobile phase A (0.1% formic acid in water) and mobile phase B (0.1% formic acid in acetontrile). The flow rate was 0.3 ml min^−1^. The linear gradient was as follows: 0–3 min: 100% A; 10–15 min: 2% A; 16–20.5 min: 100% A. The autosampler was set at 15°C. The injection volume was 1 *μ*l. Mass spectra were acquired under positive electrospray ionization with a voltage of 5,500 V. The source temperature was 450°C. The curtain gas, ion source gas 1, and ion source gas 2 were 33, 65 and 60 psi, respectively. Multiple reaction monitoring was used for quantitation with external calibration.

## Results and Discussion

### Free energies along the SMD-derived pathways do not support experimental results

To understand the increased permeability of 6DNM-NH3 relative to 6DNM through OmpF, ^8^ we first attempted to evaluate the energetics of permeation of each drug by performing independent 1D-BEUS simulations seeded by SMD simulations biasing the translational coordinate of the antibiotics along the pore axis (*Z*-axis). We performed multiple (10) SMD simulations for each drug from which an optimal permeation pathway was selected from the SMD trajectory with the lowest non-equilibrium work value (Fig. S1). A slow pulling velocity (0.5 Å ns^−1^) was employed to minimize potential hysteresis and perturbations by the drug’s induced translocation to the protein elements. The resulting pathway in each case was used to generate initial seeds for 1D-BEUS simulation for a total of 3.3 *μ*s. As expected, lack of sufficient sampling of the antibiotic orientation at the CR was apparent in the resulting 1D-BEUS simulations (Fig. S4).

Although the 1D-BEUS simulations for each drug produced apparently converging free energies (Fig. S5), a higher permeation barrier for 6DNM-NH3 was observed compared to 6DNM (Fig. S6), in disagreement with our experimental finding that 6DNM-NH3 reaches higher accumulation inside *E. coli*.^8^ This discrepancy between computation and experiment may arise because the initial pathways for BEUS, derived from SMD runs, may not have captured the most likely permeation pathway for the antibiotics. A proper assessment for the optimal initial pathway would require a more aggressive sampling of the orientation space that was not achieved in our SMD runs (Fig. S3). Although a large number of SMD runs could theoretically sample orientation space comprehensively, performing an adequate number of SMD simulations for such a complex molecular system would be computationally prohibitive.

### Free-energy along the MCPS-derived pathway supports experimental results

Since SMD-seeded BEUS simulations failed to sufficiently sample other slow DOFs such as orientations of the antibiotic (specifically at the narrow CR), we sought to first use a heuristic method to probe the multi-dimensional space more extensively, and then use this sampling to develop a better initial pathway for free energy calculations.

To enhance sampling of the orientation and translational DOFs of 6DNM and 6DNM-NH3, we developed a systematic and computationally efficient algorithm named Monte Carlo based pathway search (MCPS). MCPS determines multiple permeation trajectories through OmpF from the extracellular to periplasmic side using an energetic descriptor of the system. To run MCPS, we first generate a dataset containing hundreds of thousands of discrete drug-protein poses to exhaustively explore translational and rotational DOFs of the drug (Fig. 2A). This dataset is then used to construct a multidimensional drug-protein interaction energy (IE) landscape for the translation (*Z*-coordinate) and orientation (inclination angle, *θ*, and azimuthal angle, *ϕ*) (Fig. 2B). It is not trivial, however, to just use a rugged multi-dimensional potential energy landscape to determine the most likely pathway. ^83,84^ Additionally, the IE landscape lacks definitive start and end states for antibiotic permeation. Therefore, we used our MCPS algorithm to walk through the IE landscape using Monte Carlo (MC) moves to determine favorable trajectories connecting extracellular and periplasmic spaces (Fig. 3). The starting point within a trajectory is randomly selected from the poses closest to the extracellular space.

To comprehensively sample all possible pathways in our defined space, we obtained 2,000 MCPS trajectories that demonstrated exhaustive and convergent sampling of the conformational space (Figs. S7 and S8). Then, we used these trajectories to build a connected graph to be used in Dijkstra’s algorithm to determine the most favorable permeation pathway (Fig. 3C,D). A detailed description for each step is provided in the Methods section. Our algorithm showed an excellent computational efficiency of ≈ nearly 100 folds over 100 SMD runs (≈ 92 vs. ≈ 10K node-hours; Table 1), realizing that even this many SMD runs fail to sufficiently sample orientation space of the drug.

**Table 1:** Benchmark for calculation of most likely permeation pathway determined by our algorithm, performed on a computer with two 16 core Intel Xenon E5-2637 CPUs and one TITAN RTX GPU

| MCPS steps | Node-hour |
| --- | --- |
| Conformational search | 1.1 |
| Minimization and IE calculation | 90.5 |
| MCPS | 0.0042 |
| Most likely pathway calculation | 0.02 |
| Total: | 91.6 |

For 6DNM-NH3, the most likely pathway derived from MCPS is largely different from the one derived from the lowest-work SMD run; however, for 6DNM, the pathways derived from either approach are similar (Fig. S12). The MCPS-derived pathways were then used as initial seeds for two independent 1D-BEUS simulations for either drug to obtain free-energy profiles.

The free-energy profiles revealed that within the CR, both drugs face high permeation energetic barriers, similar to the SMD-derived 1D-BEUS; however, the overall free-energy barrier for 6DNM-NH3 is significantly less than 6DNM (by 2 kcal mol^−1^), (Fig. 4A). The barrier arises likely because: (i) the narrowness of the CR causes a loss in entropy, which has been shown to contribute to the free-energy barrier for other antibiotics’ translocations through OmpF in previous MD simulation studies; ^20^ and (ii) the loss of drug-water interactions at the CR leads to a large desolvation enthalpic penalty (Fig. 4C), which has been shown to contribute to the overall free-energy barrier in several biological processes such as protein folding, ^85^ and protein-ligand binding. ^86^ However, the lower free-energy barrier for 6DNM-NH3 can be attributed to the increased protein-drug interactions enabled by the positively charged amino group (amine1) (Fig. 4B,C), which offers more enthalpic compensation to the entropic and desolvation costs within the CR for the aminated compound.

**Figure 4:**
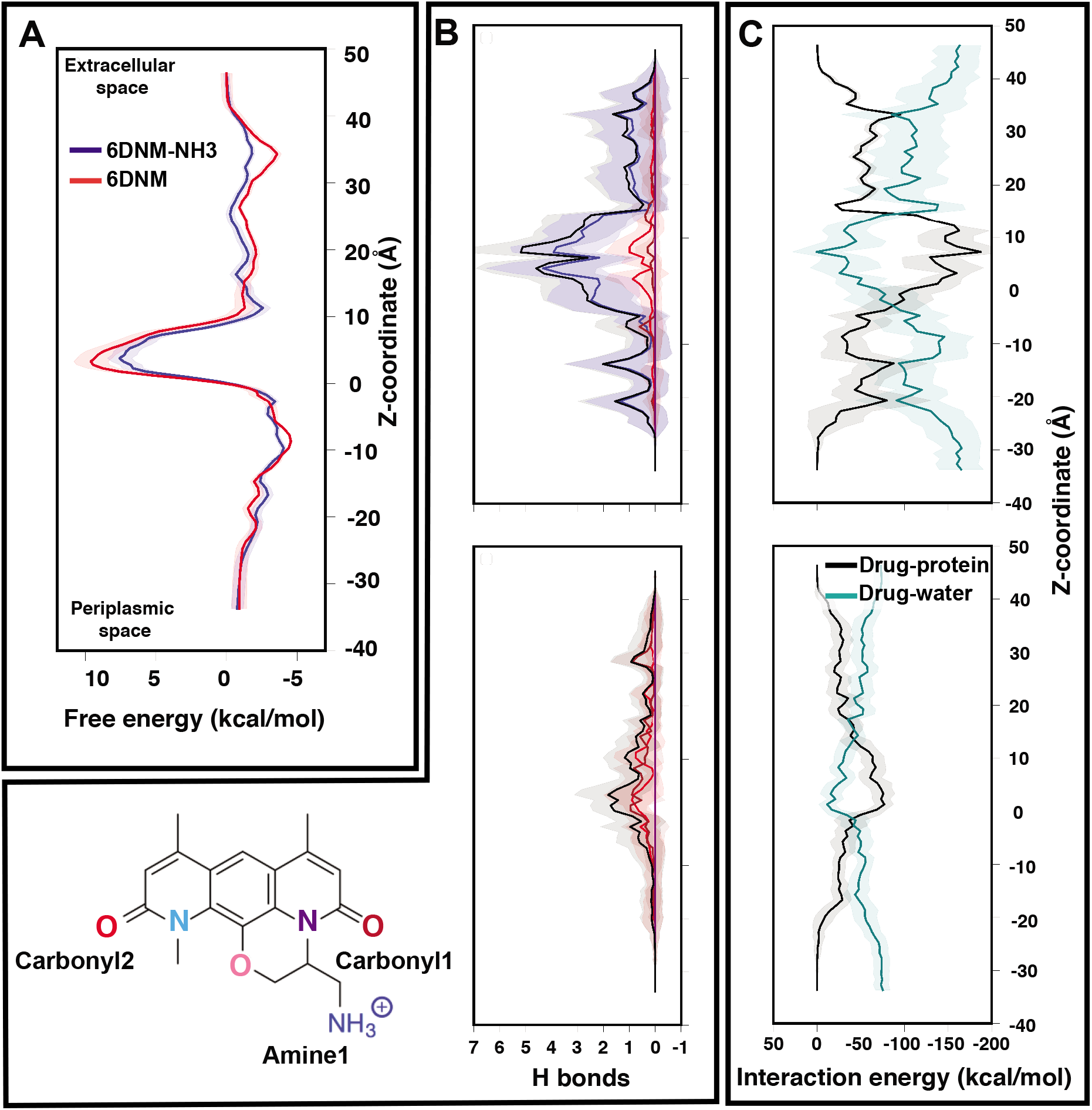
Comparison of the energetics of permeation for 6DNM-NH3 and 6DNM through OmpF. (A) Mean (solid) and standard deviation (shaded) of the free-energy for permeation of 6DNM-NH3 (blue) and 6DNM (red). Free energies were calculated using all the MCPS-BEUS windows and projected along the *Z*-coordinate of the antibiotic C.O.M. The location of the CR is highlighted in orange. (B) Mean and standard deviation of the number of hydrogen bonds between any protein residue and each functional group in the drug (colored differently according to chemical structure shown in bottom, left) of 6DNM-NH3 (*top*) and 6DNM (*bottom, right*) at different *Z*-positions. The total number of hydrogen bonds for each drug is colored in black. The nomenclature and color code for each electronegative atom are shown in the bottom left panel. (C) Mean and standard deviation of the drug-protein and drug-water interaction energy at different *Z*-positions for 6DNM-NH3 (*top*) and 6DNM (*bottom*).

We then used the MCPS-BEUS free-energy profiles combined with our computed diffusion coefficients along the pore axis to determine permeability of either drug through OmpF. Our calculations showed a 30-fold greater permeability for the aminated drug (1.1 × 10^−3^ cm/s for 6DNM-NH3 vs. 3.3 × 10^−5^ cm/s for 6DNM), substantiating the increased accumulation of 6DNM-NH3 into *E. coli* we observed experimentally. ^8^

Notably, for 6DNM-NH3, a significant decrease (by ≈ 3 kcal/mol) in the energetic barrier was observed in the free-energy profile derived using MCPS-BEUS compared to the one derived using SMD-BEUS. For 6DNM, the barrier calculated by the two approaches are rather similar, possibly because the initial pathways derived from these approaches are similar (Figs. S11 and S12).

### Permeation mechanism of 6DNM-NH3 through OmpF

To understand the mechanisms by which the primary amine group (amine1; Fig. 4) enhances the permeability of 6DNM-NH3, we determined the most likely permeation pathway through the newly generated free-energy landscape for each drug. As we are mainly interested in the translocation of the antibiotics through the CR, we performed further analysis of the frames within the entrance, top, bottom, and exit of the CR along the resulting pathways. At each location, *θ*, *ϕ* and hydrogen bonding pattern with protein residues were analyzed (Figs. 5 and 6).

**Figure 5:**
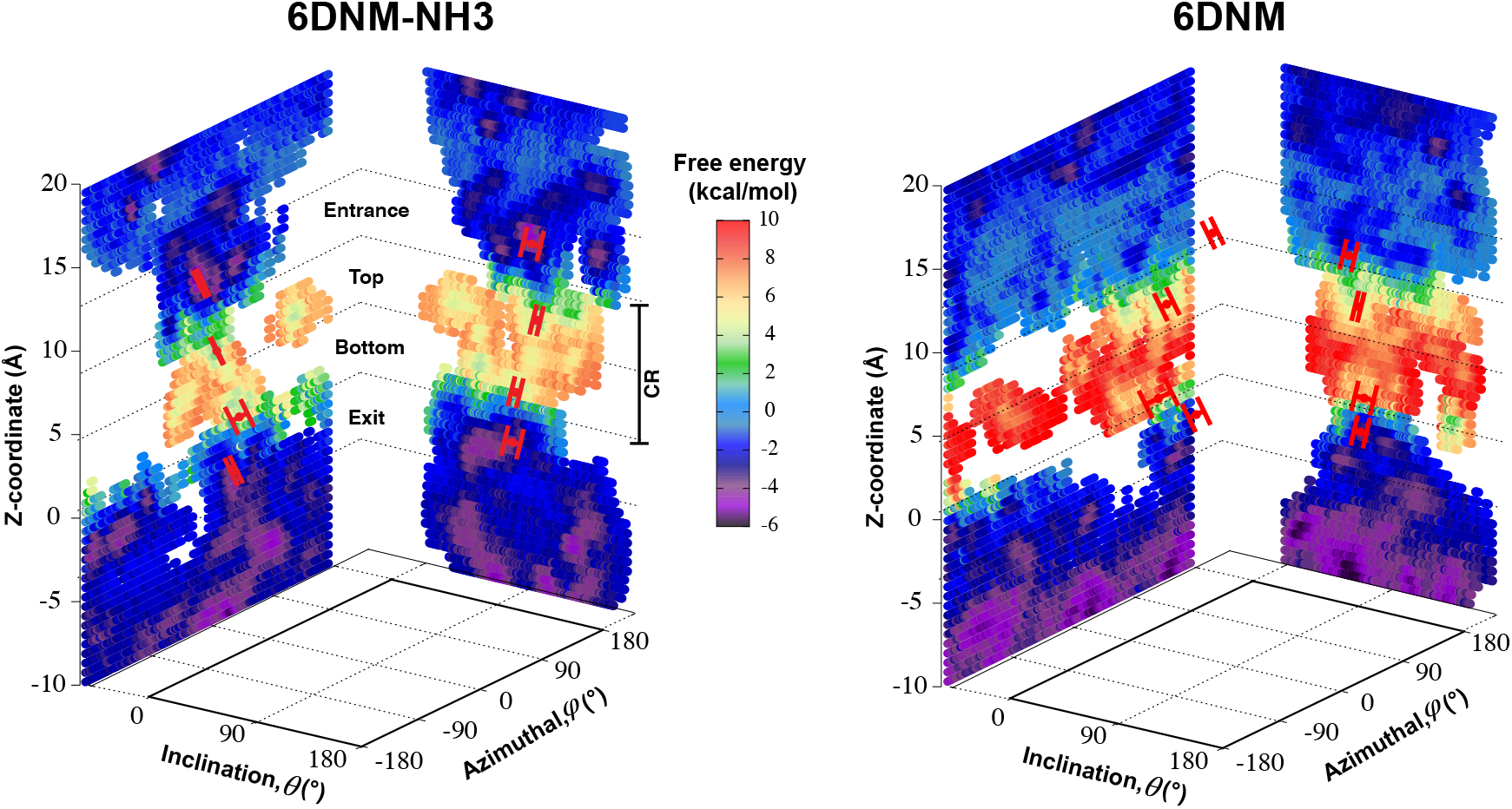
Mechanism of antibiotic permeation through OmpF. Free-energy landscapes for permeation of 6DNM-NH3 and 6DNM (calculated using all respective MCPS-BEUS windows), projected along the *Z*-coordinate, inclination (*θ*) and azimuthal (*ϕ*) angle of the antibiotic. All free-energy values were calculated relative to the free energy in bulk solution. Mean and standard deviation of *θ* and *ϕ* of the antibiotic in the CR at 1) the entrance (*Z* = 9–12 Å), 2) top (*Z* = 5–8 Å), 3) bottom (*Z* = 1–4 Å), and 4) exit (*Z* = – 3–0 Å) of the CR, along the most likely pathway are highlighted in red.

**Figure 6:**
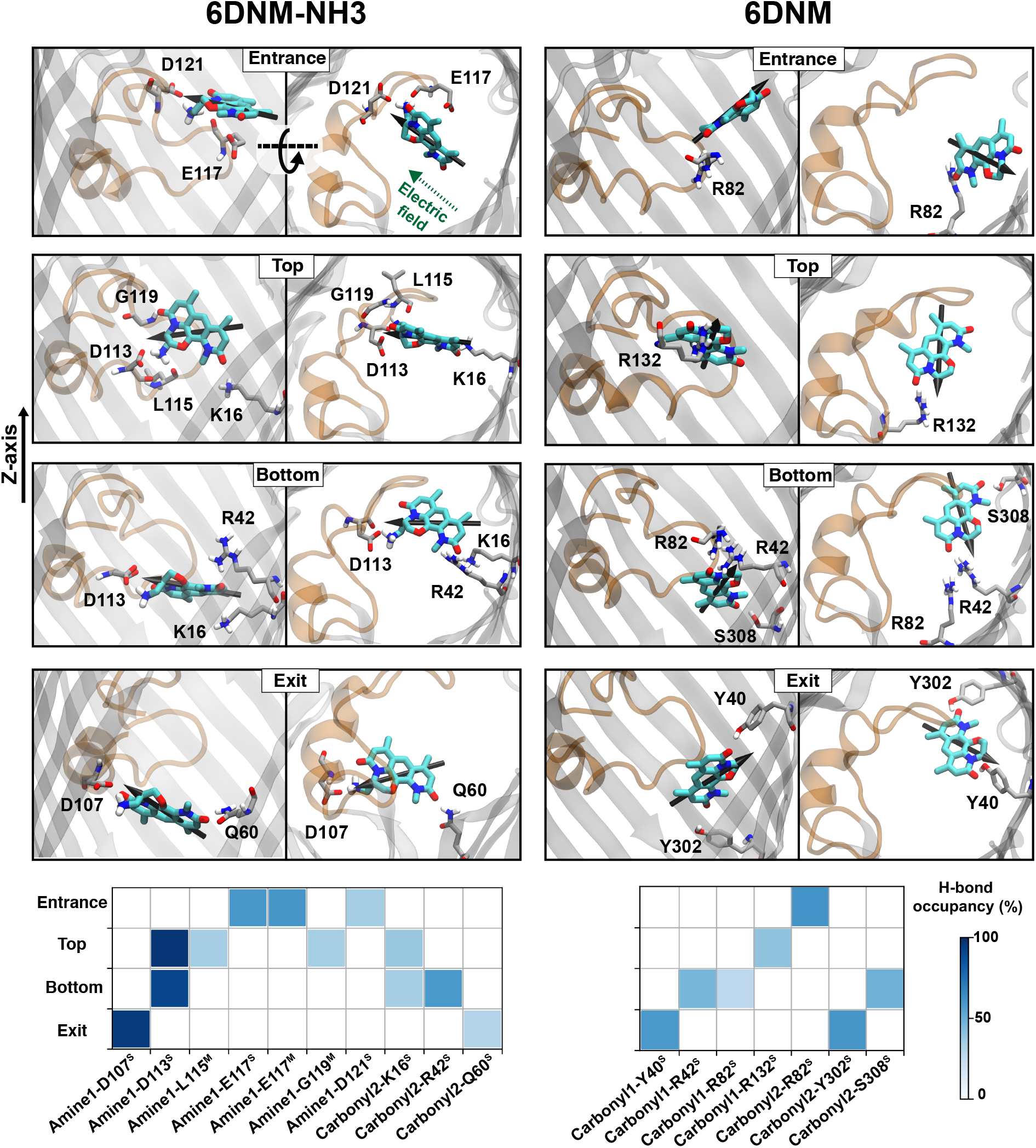
Key interactions along the antibiotic permeation pathway. (*Top*) Snapshots of both drugs at different locations along the CR, highlighting key interacting residues. (*Bottom*) Hydrogen bonding between the different functional groups of 6DNM-NH3 and 6DNM and protein elements at different locations along the CR. The interactions shown have hydrogen bond probability greater than 20%. Nomenclature for the antibiotic functional groups is provided in Fig. 4. Superscripts S and M represent side- and main-chain of protein residues, respectively.

We find that upon entering the CR, 6DNM-NH3 orients such that its vector (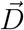, approximately aligned with the drug’s dipole moment with a magnitude of 19 Debye), aligns with the transverse electric field in the CR (*θ* = 94.0 ± 9.2° and *ϕ* = −5.1 ± 7.7°) and maintains this orientation throughout the entire CR (from entrance to exit, Fig. 5). The observed alignment of the dipole of 6DNM-NH3 with the protein’s electric field may compensate the reduced entropy of the drug within the CR, as illustrated by the appearance of a local affinity site at the CR entrance (Fig. 5). This affinity site helps to stabilize 6DNM-NH3 within the CR, which could increase the probability of the drug passing through the porin. Accordingly, previous MD simulation studies have suggested that electric field alignment of the dipole of other antibiotics and the resulting affinity sites to be beneficial in permeation through OmpF.^20,24,25,27^ In contrast, for 6DNM, no such local affinity site was observed at the CR entrance since the dipole moment of 6DNM is significantly weaker (8 Debye) than that of 6DNM-NH3 (Fig. 5). As a result, throughout the CR, the free energy for 6DNM in both projections is calculated to be significantly greater than that of 6DNM-NH3 (Fig. 5).

The particular orientation of 6DNM-NH3 within the CR also poses amine1 to interact with acidic residues of L3 and carbonyl2 to interact with residues of the basic ladder located at the barrel wall (Figs. 5 and 6). At the CR entrance, amine1 interacts with the side chain of D121 and side/main chain of E117, located near the extracellular terminal of L3; another factor (in addition to dipole alignment) contributing to the appearance of a local affinity site at this location (Fig. 5). Within the CR, however, amine1 breaks these interactions to interact with the side chain of D113 and main chains of L115 and G119. At this location, carbonyl2 also interacts with the side chains of K16 and R42, further stabilizing the drug in the CR. Notably, the downward movement of the antibiotic towards the periplasmic space is accompanied with a drug rotation (monitored by 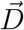), while amine1 and carbonyl2 maintain similar interactions along the pore axis; this rotation is expected to help the bulky body of the drug to advance through the narrow CR. The drug exits the CR with no further reorientation; however, interactions are broken as amine1 and carbonyl2 interact with the side chains of D107 and Q60, respectively. Overall, our analysis shows that 6DNM-NH3 permeates through the CR via a stepwise mechanism as the drug hops in order for amine1 and carbonyl2 to interact with residues along the pore. In contrast, for 6DNM, significantly fewer hydrogen bonds and a completely different orientation was observed because this antibiotic lacks a primary amine and carries a weaker dipole moment (Figs. 5 and 6).

Our mechanism for permeation of 6DNM-NH3 through OmpF could be generalized to other general-diffusion porins of bacteria, such as OmpC (*E. Coli*), OmpK36 (*K. pneumoniae*) and OmpE36 (*E. cloacae*), since these porins have a similar architecture to OmpF and conserve several key residues involved in the permeation process (Fig. S13).

### D113 mutation decreases 6DNM-NH3 accumulation inside *E. coli*

Our computational analysis showed that amine1-D113 is the most persistent interaction along the entire CR (Figs. 5 and 6). To validate the importance of this interaction in permeation of 6DNM-NH3 through OmpF, we compared experimentally the accumulation of 6DNM-NH3 in *E. coli* BW25113 (parent strain) with two strains containing single mutants in OmpF, *E. coli* OmpF^D113V^ and *E. coli* OmpF^D113F^. Based on our simulation results, we expect cells expressing D113V- and D113F-OmpF to accumulate 6DNM-NH3 to a lesser extent than cells expressing WT-OmpF, because substitution of D113 with uncharged amino acids can disrupt favorable interactions with amine1 at the CR, which would enthalpically compensate for the entropic loss of the drug in this region. The loss of this favorable interaction is therefore expected to decrease the rate of permeation through the porin in the designed OmpF mutants. In accordance with our expectations, we observed a 2.5 fold decrease in the accumulation for cells expressing either mutant (Fig. 7).

**Figure 7:**
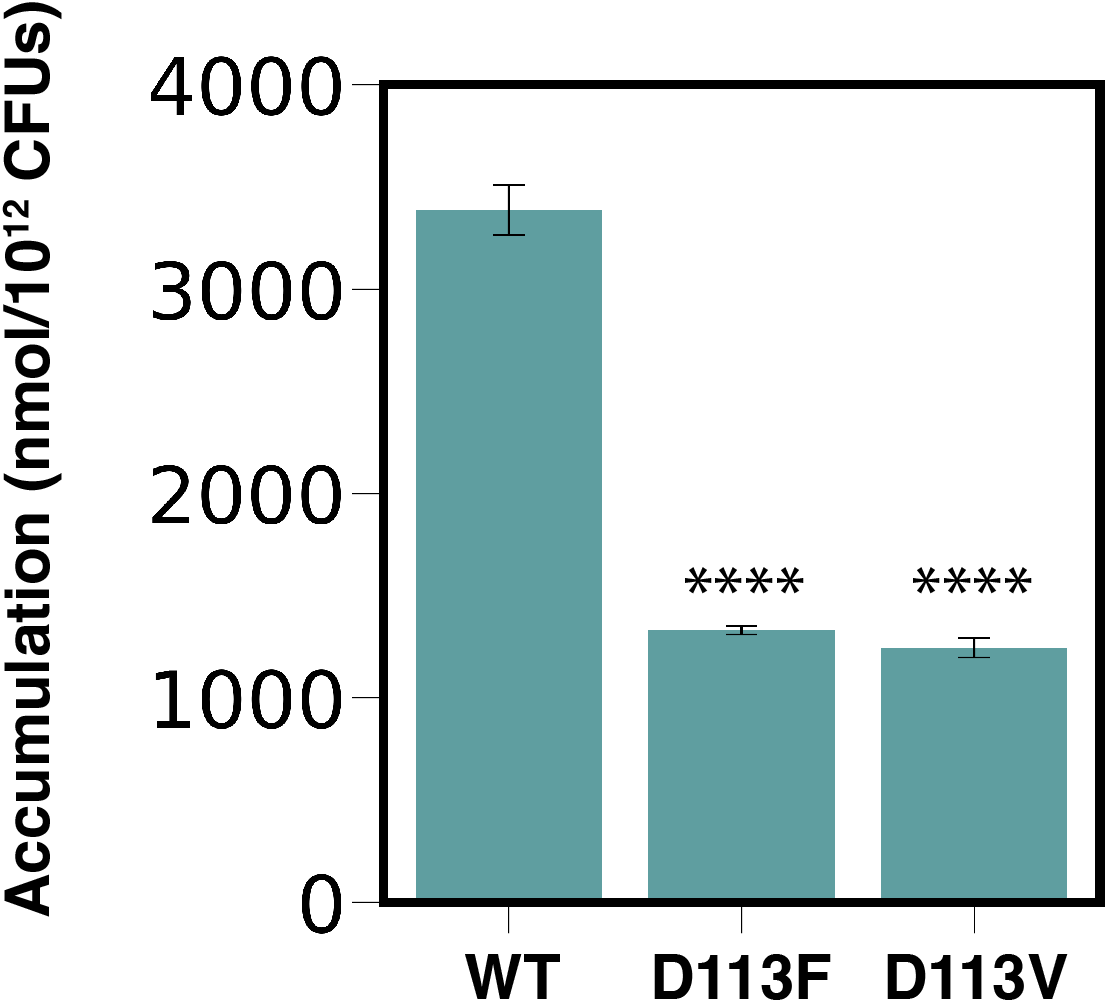
6DNM-NH3 accumulation in *E. coli* BW25113 (WT) and two strains containing single mutants in OmpF: *E. coli* OmpF^D113V^ and *E. coli* OmpF^D113F^. Accumulation reported in nmol per 10^12^ colony-forming units (CFUs). Data shown represent the average of three independent experiments. Error bars represent the standard deviation of the data. Statistical significance was determined by a two-sample Welch’s t-test (one-tailed test, assuming unequal variance) relative to accumulation in the WT strain. Statistical significance is indicated with asterisks (**** p < 0.0001).

## Concluding Remarks

MD simulations can provide important structural/dynamic information on molecular processes that can not be obtained using typical experimental methods due to their inherently limited temporal/spatial resolutions. However, despite the ever-increasing capability of supercomputers, the typical timescale of atomistic MD simulations is insufficient to adequately sample configuration space for slow molecular processes. This prevents an accurate description of processes such as membrane transport which occur on long timescales (milliseconds or longer) and include slow degrees of freedoms (DOFs). To alleviate this problem enhanced sampling methods that can bias sampling along the most relevant DOFs are used; however, sufficient sampling of configuration space still remains daunting unless additional simplifications are made.

One such simplification is to focus the calculations on a pathway that describes the most relevant DOFs during a structural transition. Identifying this transition pathway using conventional methods can still be computationally expensive; therefore, we have developed an efficient, heuristic approach using a combination of Monte Carlo simulation and graph theory. Within this approach, we first create a dataset of discrete energy minimized molecular states using a grid-based workflow.^51^ This dataset could be created for any system by exploring the states along all the relevant DOFs and approximately calculating the energy associated withe each state. We then use the resulting dataset to determine multiple energetically favorable trajectories along the molecular process of interest using our novel Monte Carlo based pathway search (MCPS) algorithm. Finally, we determine the most likely pathway sampled in our MCPS trajectories using Dijkstra’s algorithm and use this pathway to determine the free-energy underlying the molecular process with established enhanced sampling techniques such as BEUS.

We applied our approach to investigate antibiotic permeation through outer membrane (OM) porins of Gram-negative bacteria; a process that involves multiple slow DOFs, most importantly translation and orientation of the drug. This application is highly biomedically relevant since these pathogens are becoming increasingly resistant to currently available antibiotics, and the development of new antibiotics targeting them has been slow.^1,2^ Specifically, we investigated the molecular basis for the conversion of Gram-positive-only to broad-spectrum antibiotics with the addition of a primary amine group as observed in our previous experiments. We found that the free-energy barrier for permeation was significantly lower for the aminated derivative corresponding to a greater permeability through OM porins. Further analysis revealed that the added amino group facilitates favorable antibiotic permeation through porins by enabling the drug molecule to align its dipole to the intrinsic electric field of the porin and by forming interactions with several charged residues. The importance of these interactions, which appear to be conserved among different species, to effective permeation across the outer membrane of Gram-negative bacteria was further validated with experimental mutagenesis and whole-cell accumulation assays. In general, the structural insights offered by this study on the mechanisms by which the primary amine enhances permeation of antibiotics could help to convert other Gram-positive-only antibiotics into broad-spectrum antibiotics. Furthermore, our novel computational method could be generalized to sample better other long-timescale molecular processes involving slow DOFs.

## Acknowledgements

This research is supported by National Institutes of Health grants R01-AI136773 (to PH and ET), R01-HL131673 (to ET), and P41-GM104601 (to ET). Simulations in this study have been performed using allocations at National Science Foundation Supercomputing Centers (XSEDE grant number MCA06N060 to ET) and the Blue Waters supercomputer of National Center for Supercomputing Applications at University of Illinois at Urbana-Champaign.

