## Supplementary Information for "Rationalizing generation of broad spectrum antibiotics with the addition of a primary amine"

### Supporting Information

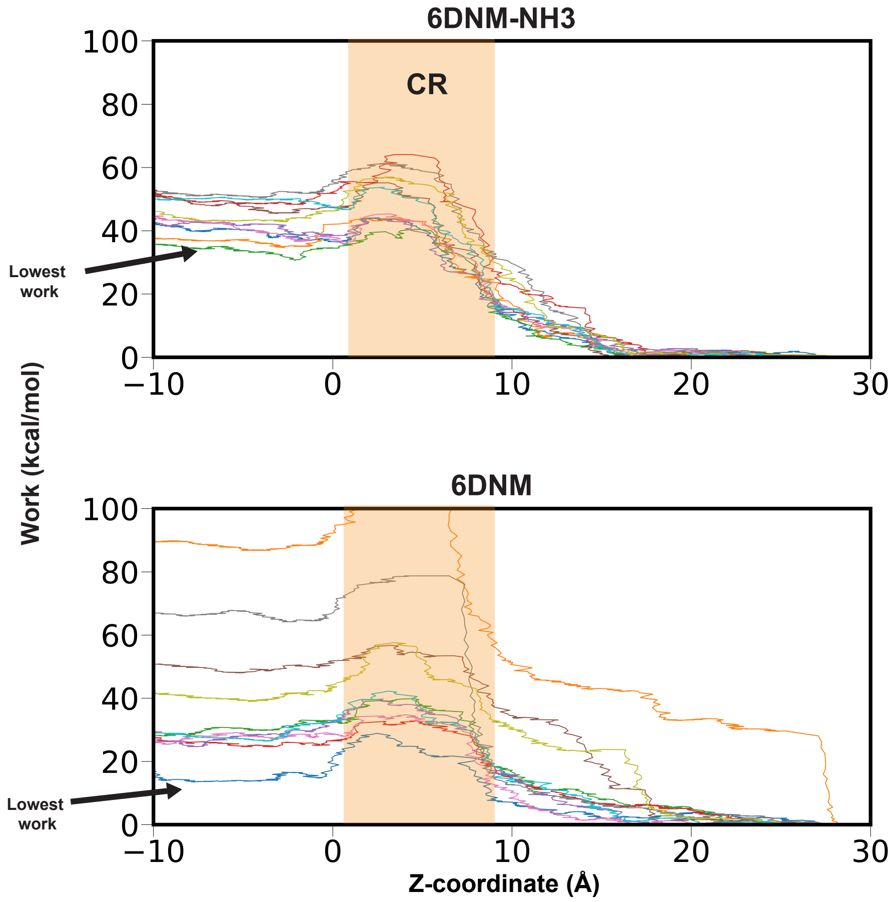


Figure S1: The accumulated non-equilibrium work during different independent SMD simulations (represented in unique color) of 6DNM-NH3 (*top*) and 6DNM (*bottom*). Work values are projected onto the position of the drug along the membrane normal ($z$-axis) and relative to the midplane of the membrane ($z = 0$). The location of the CR is highlighted in orange.

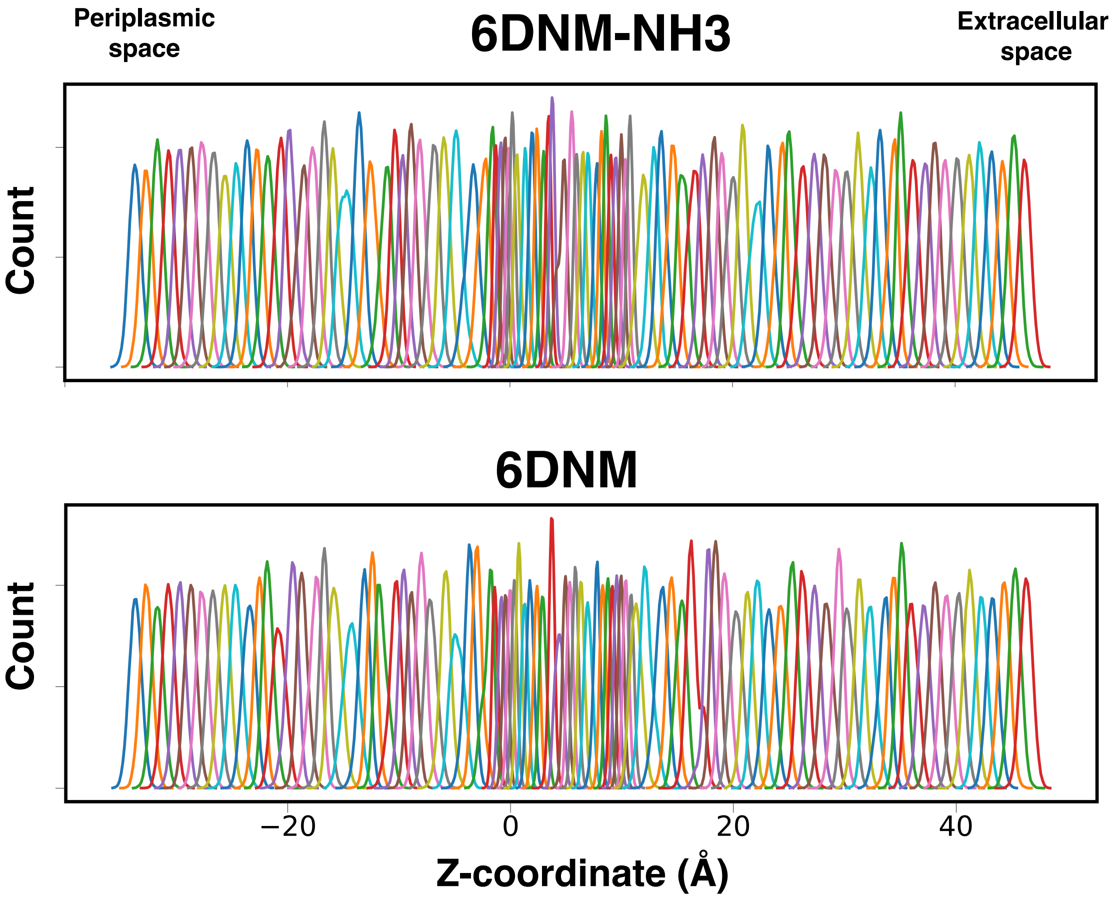


Figure S2: Window overlaps from SMD-seeded BEUS simulations for permeation of 6DNM-NH3 (*top*) and 6DNM (*bottom*) through OmpF. The histograms show distribution probabilities of the drug along the membrane normal ($Z$-axis) in each replica (window). Windows are spaced at 1 Å intervals for a span of 80 Å extending from the periplasmic ($Z = -34$ Å) to the extracellular bulk solution ($Z = 46$ Å), except for the region between the entrance and exit of the CR ($Z = -3$ to 12 Å) where a 0.5-Å spacing was used to ensure adequate histogram overlap.

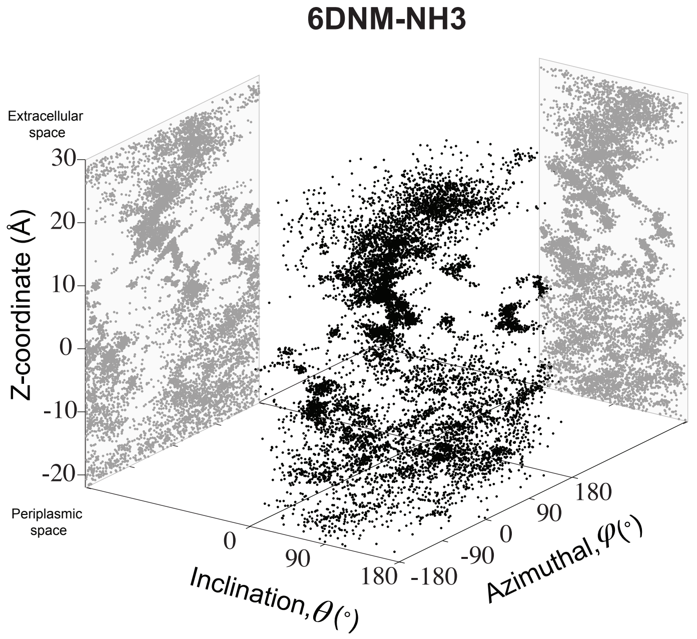


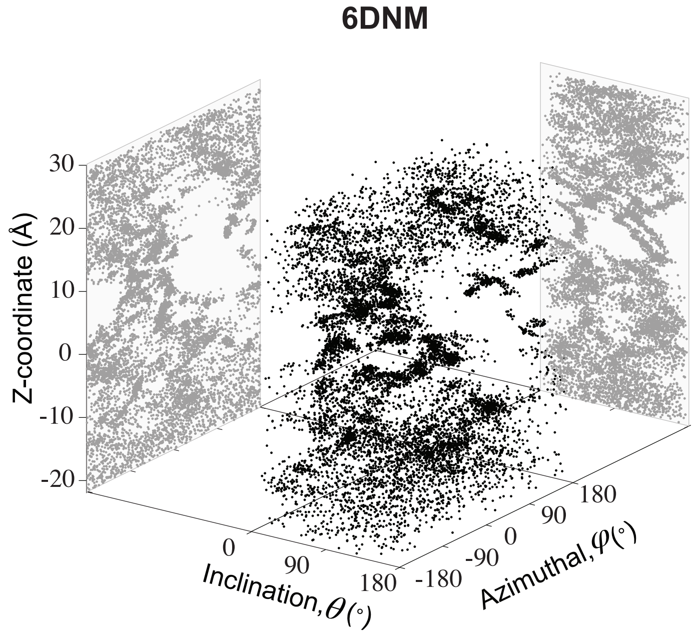


Figure S3: Sampling of 6DNM-NH3 (*top*) and 6DNM (*bottom*) during their respective 10 independent SMD simulations projected on the orientation (inclination and azimuthal) and translation ($Z$-coordinate) DOFs of the antibiotic.

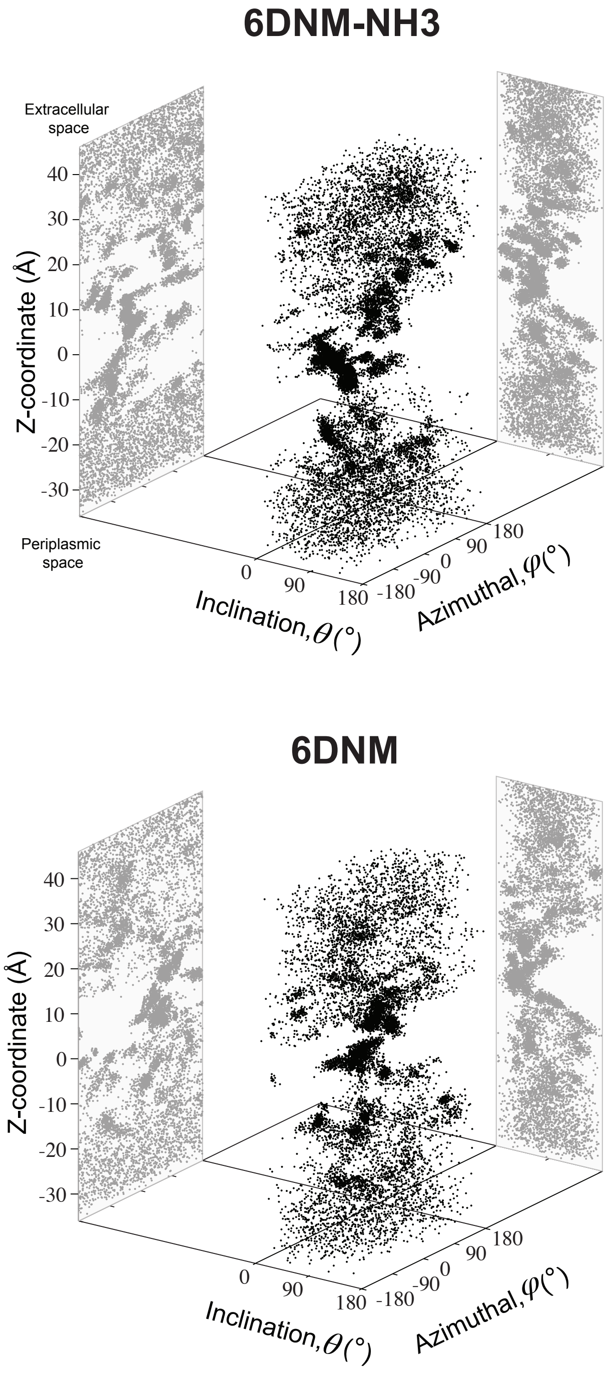


Figure S4: Sampling of 6DNM-NH3 (*top*) and 6DNM (*bottom*) during their respective SMD-seeded BEUS simulations projected on the orientation (inclination and azimuthal) and translation ($Z$-coordinate) DOFs of the antibiotic.

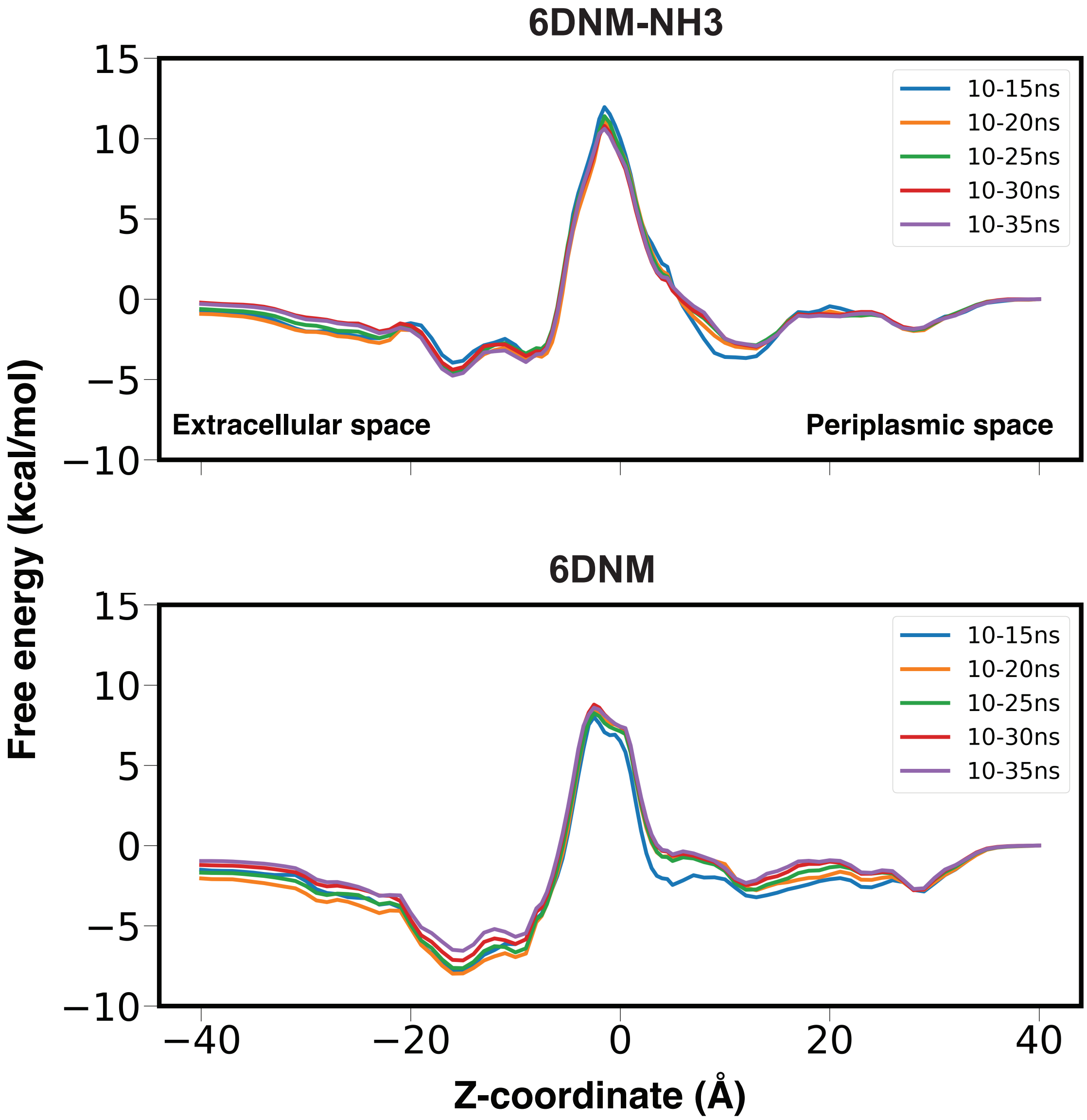


Figure S5: Convergence of potential of mean force (PMF) calculations derived from SMD-seeded BEUS was monitored by calculating permeation free energy for 6DNM-NH3 (*top*) and 6DNM (*bottom*) for different simulation segments in each BEUS window after discarding the first 10 ns (10–15 ns, 10–20 ns, 10–25 ns, 10–30 ns, and 10–35 ns). Free energies are projected onto the $Z$-coordinate of the C.O.M of each drug.

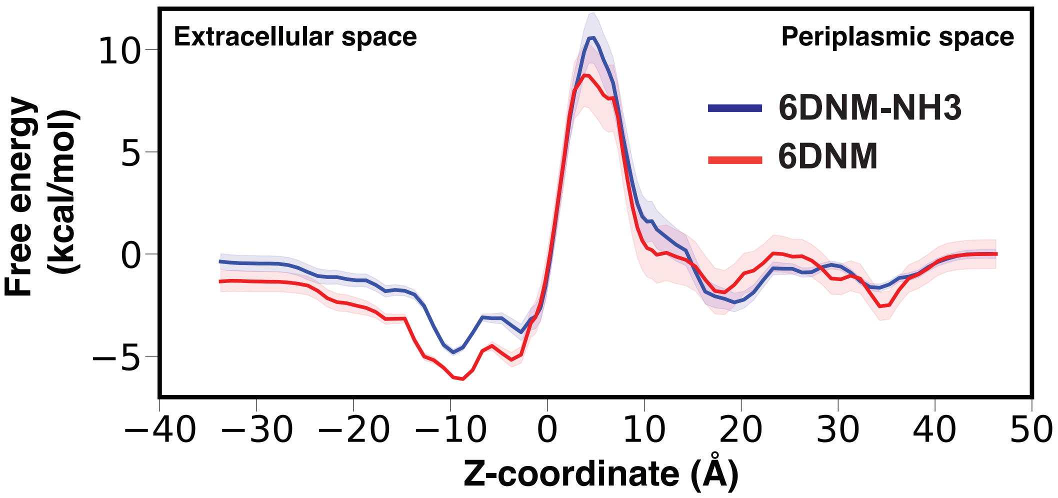


Figure S6: Mean (solid) and standard deviation (shaded) of the free energy values for the permeation of 6DNM-NH3 (blue) and 6DNM (red), calculated using all the SMD-seeded BEUS windows, projected along the $Z$-coordinate of the antibiotic C.O.M.

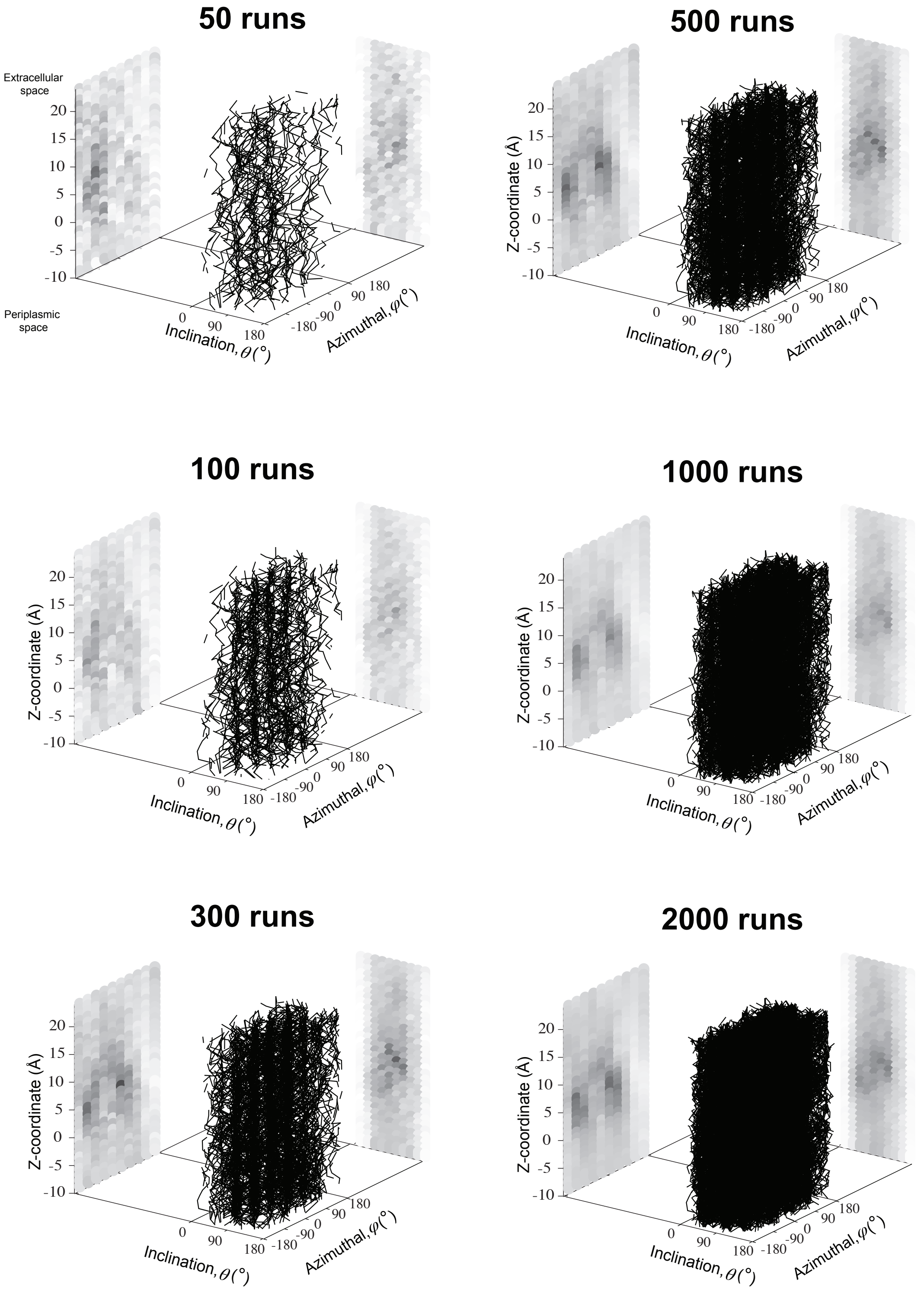


Figure S7: Convergence of MCPS runs of 6DNM-NH3, determined by projecting the Boltzmann weighted densities of the MCPS trajectories at different numbers of runs along the $Z$-coordinate, inclination ($\theta$) and azimuthal ($\phi$) of the antibiotic.

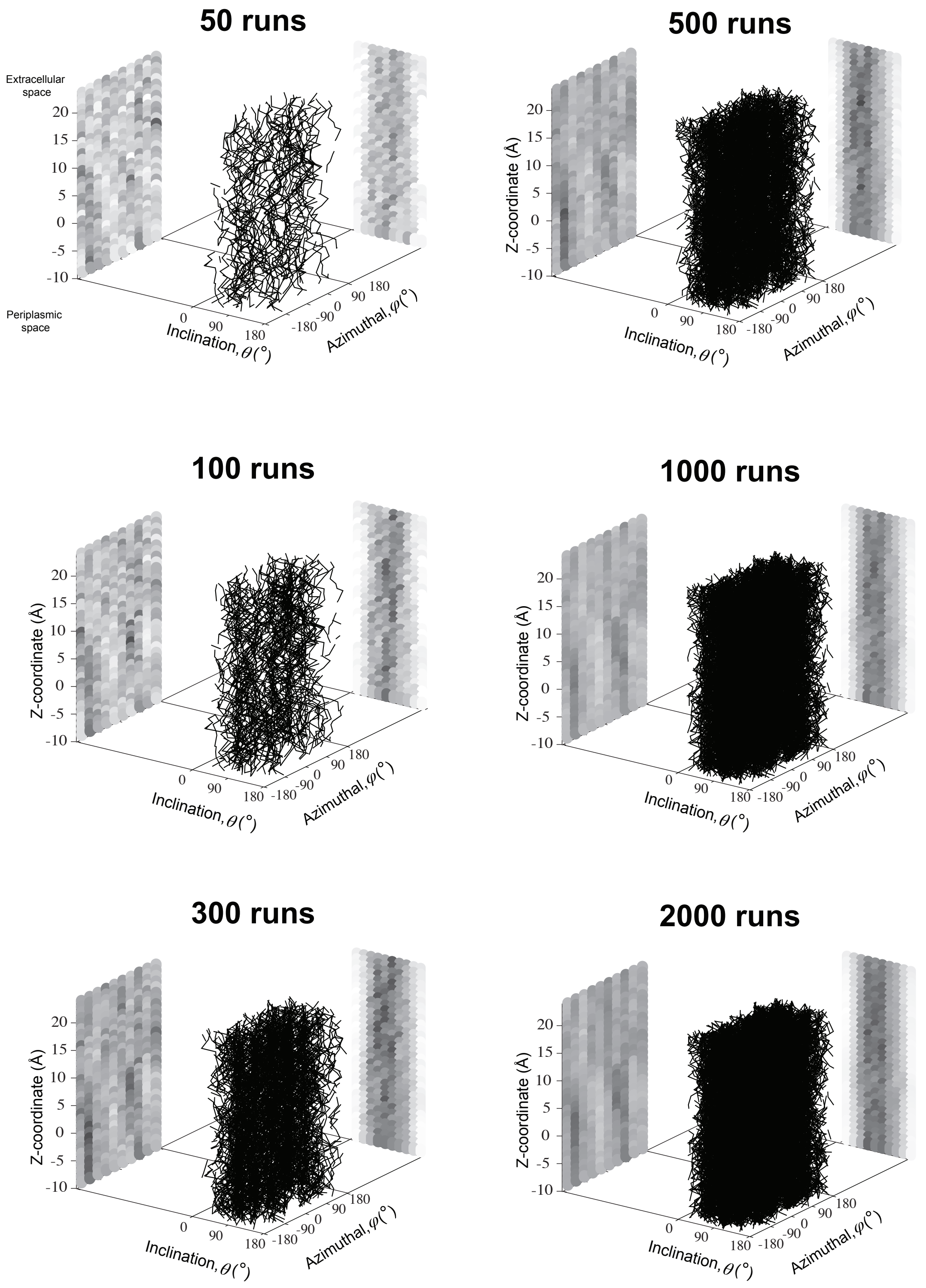


Figure S8: Convergence of MCPS runs of 6DNM, determined by projecting the Boltzmann weighted densities of the MCPS trajectories at different number of runs along the $Z$-coordinate, inclination ($\theta$) and azimuthal ($\phi$) of the antibiotic.

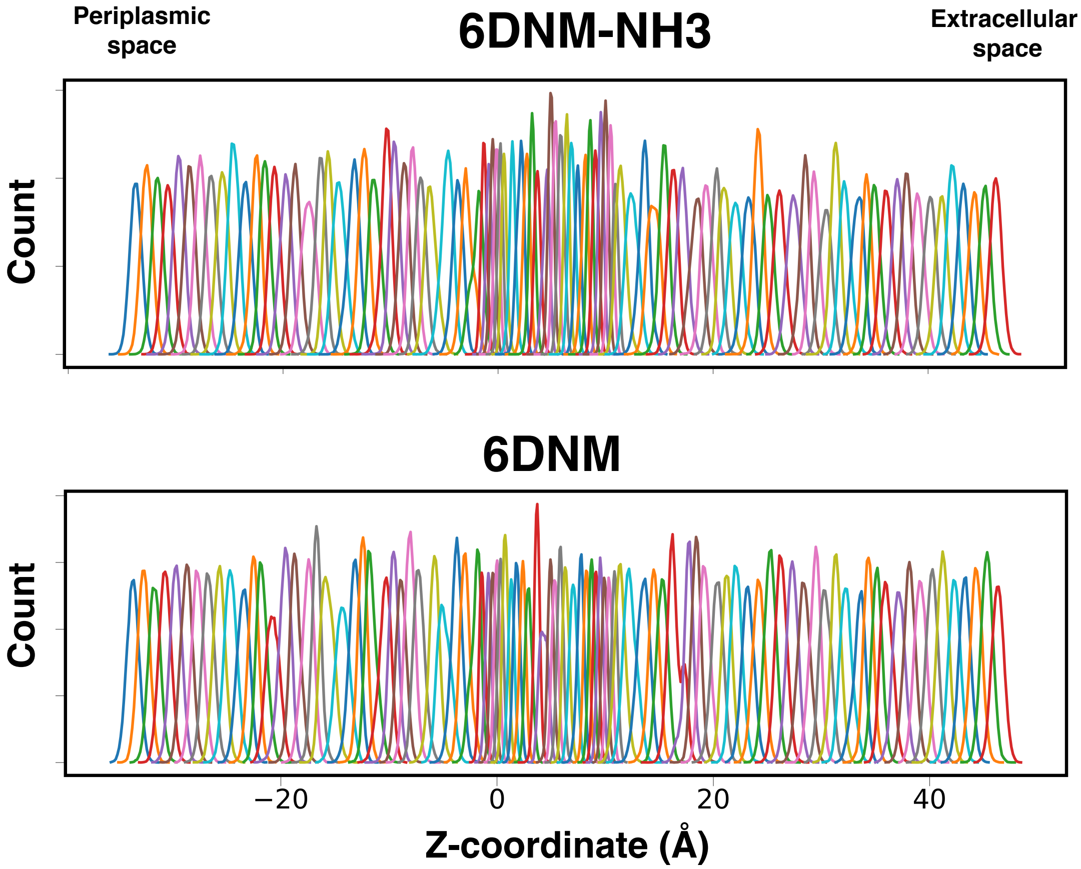


Figure S9: Window overlaps from MCPS-seeded BEUS simulations for permeation of 6DNM-NH3 (*top*) and 6DNM (*bottom*) through OmpF. The histograms show distribution probabilities of the drug along the membrane normal ($Z$-axis) in each replica (window). Windows are spaced at 1 Å intervals for a span of 80 Å extending from the periplasmic ($Z = -34$ Å) to the extracellular bulk solution ($Z = 46$ Å), except for the region between the entrance and exit of the CR ($Z = -3$ to 12 Å) where a 0.5-Å spacing was used to ensure adequate histogram overlap.

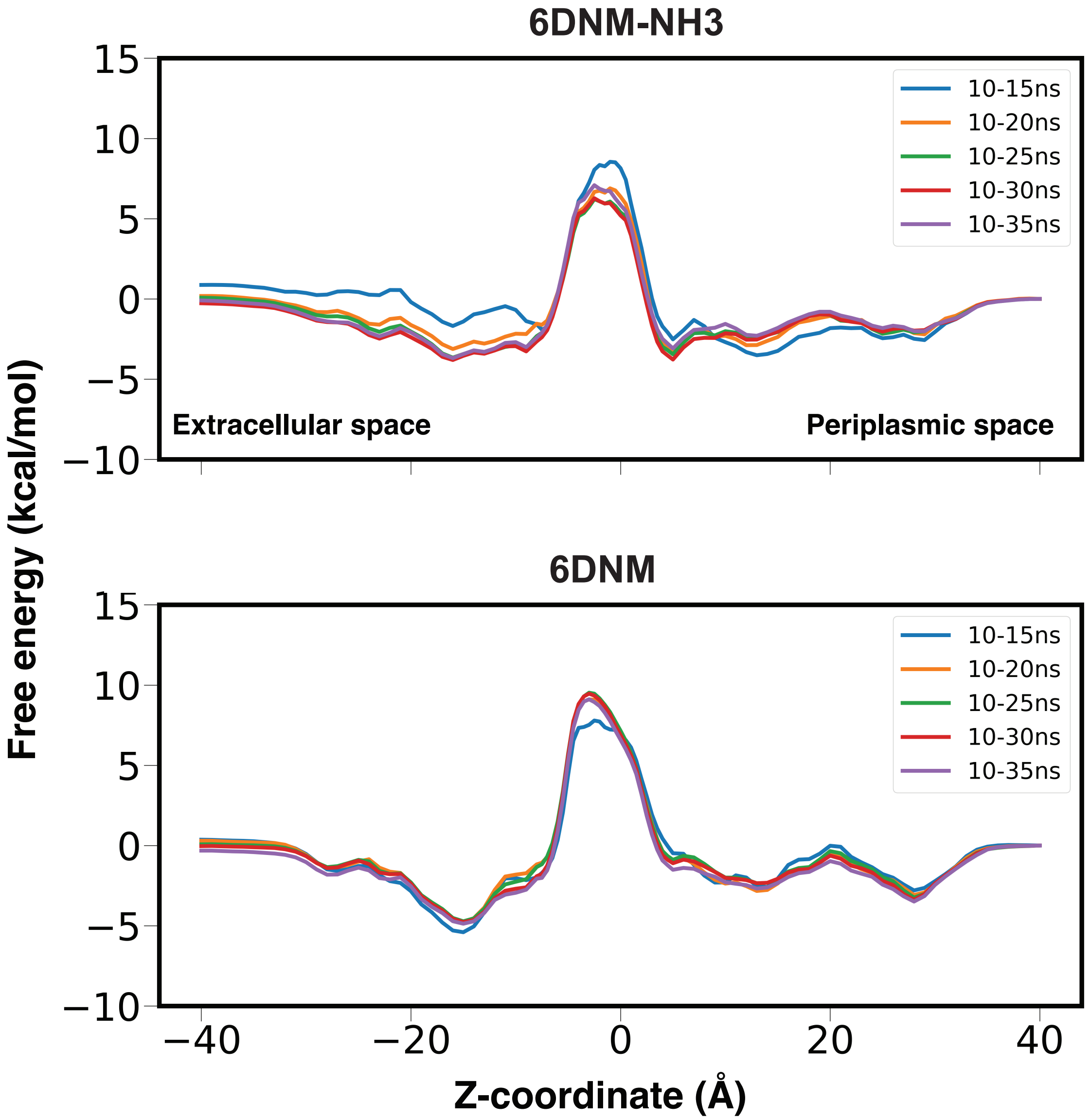


Figure S10: Convergence of PMF calculations for MCPS-seeded BEUS simulations was monitored by calculating permeation free energy for 6DNM-NH3 (*top*) and 6DNM (*bottom*) at different simulation times in each BEUS window after discarding the first 10 ns (10–15 ns, 10–20 ns, 10–25 ns, 10–30 ns, and 10–35 ns). Free energies are projected onto the $Z$-coordinate of the C.O.M of the drug.

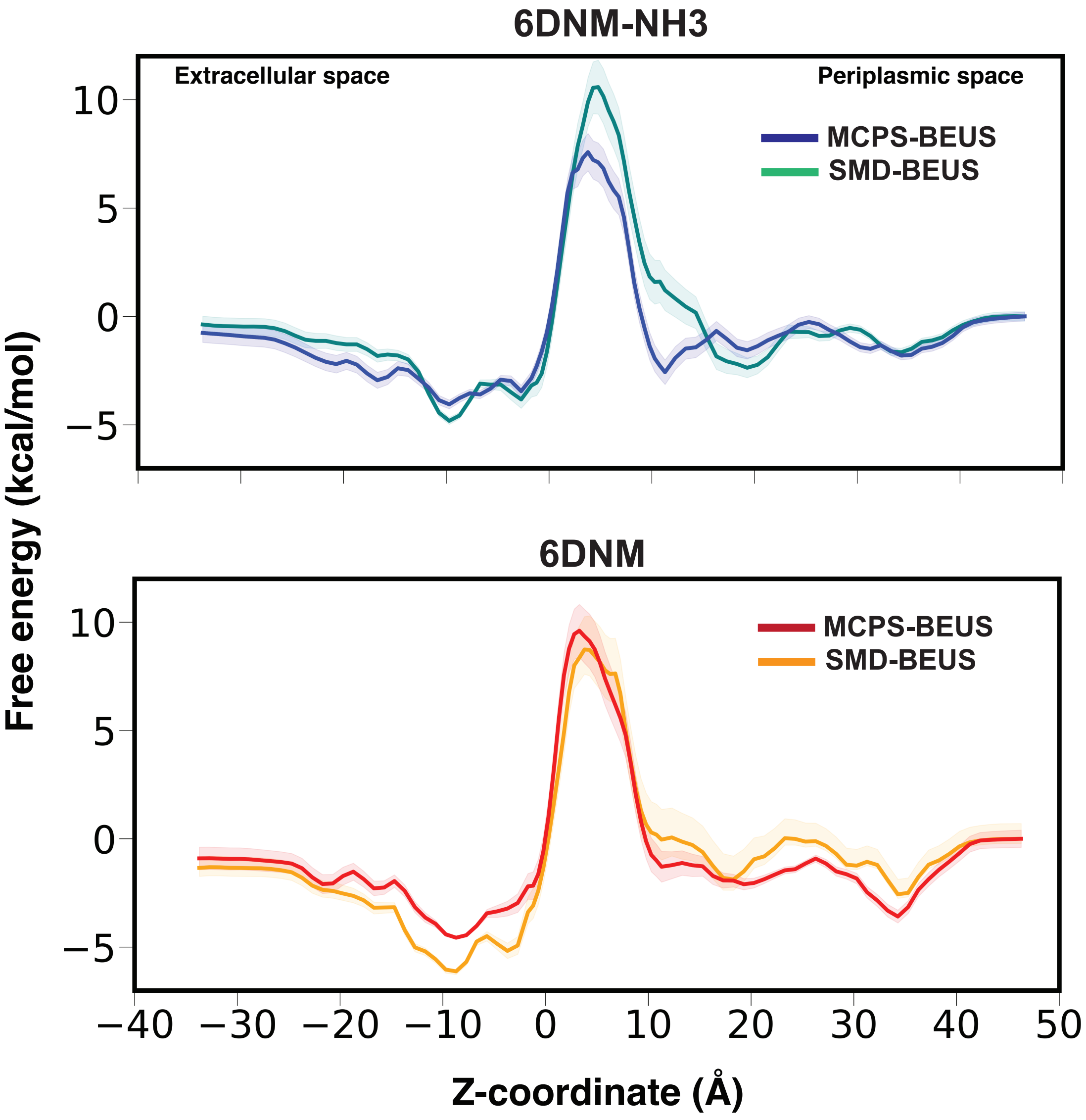


Figure S11: Comparison of free energy of permeation derived from SMD-seeded BEUS and MCPS-seeded BEUS simulations for 6DNM-NH3 (*top*) and 6DNM (*bottom*). For 6DNM-NH3, a significant decrease (by ≈3 kcal/mol) in the energetic barrier is observed in the free energy profile derived using MCPS-seeded BEUS than the one derived from SMD-seeded BEUS. For 6DNM, the barrier remains similar in both approaches.

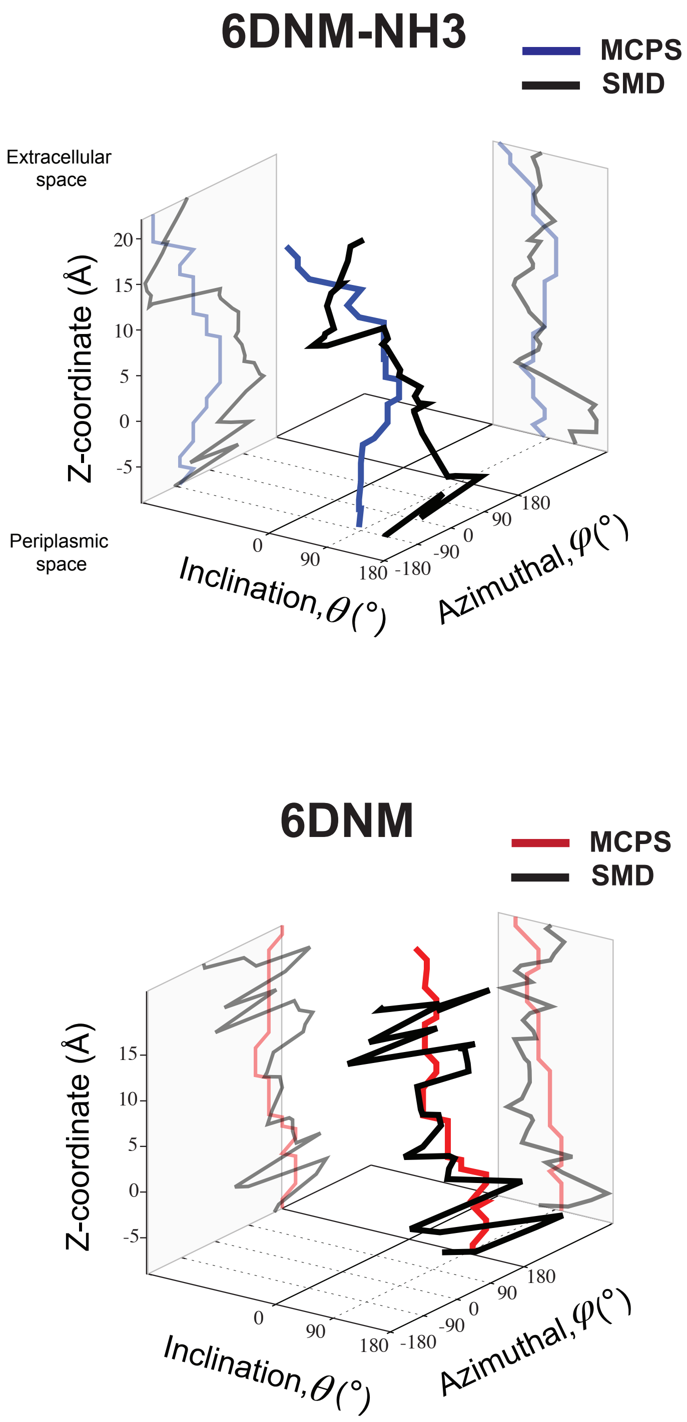


Figure S12: Comparison of the initial permeation pathways derived using MCPS and SMD for 6DNM-NH3 (*top, blue*) and 6DNM (*bottom, red*) through OmpF. Pathways are projected onto the orientation (inclination and azimuthal) and $Z$-coordinate of the antibiotic. These pathways were used to seed the BEUS simulations.

```
                16                       42                 60
OmpF:   AEIYNKDGNKVDLYGKAVGLHYFSKGNGENSYGGNGDMTYARLGFKGETQINSDLTGYGQ 60
OmpC:   AEVYNKDGNKLDLYGKVDGLHYFSDNKD-----VDGDQTYMRLGFKGETQVTDQLTGYGQ 55
OmpE36: AEIYNKDGNKLDLYGKAVGLHYFSDNDG-----NDGDKTYARLGFKGETKINDQLTGYGQ 55
OmpK36: AEIYNKDGNKLDLYGKIDGLHYFSDDKS-----VDGDQTYMRVGVKGETQINDQLTGYGQ 55

                                                107 113 115 117 119
OmpF:   WEYNFQGNNSEGADAQTGNKTRLAFAGLKYADVGSFDYGRNYGVVYDALGYTDMLPEFGG 120
OmpC:   WEYQIQGNSAEN-ENN--SWTRVAFAGLKFQDVGSFDYGRNYGVVYDVTSWTDVLPEFGG 112
OmpE36: WEYNFQGNNSEGADAQSGNKTRLAFAGLKFGDAGSFDYGRNYGLVYDAIGITDMLPEFGG 115
OmpK36: WEYNVQANNTESSSDQ--AWTRLAFAGLKFGDAGSFDYGRNYGVVYDVTSWTDVLPEFGG 113
                                                L3 residues

        121
OmpF:   DTAY-SDDFFVGRVGGVATYRNSNFFGLVDGLNFAVQYLGKNER--------------DT 165
OmpC:   DTYG-SDNFMQQRGNGFATYRNTDFFGLVDGLNFAVQYQGKNGNPSGEGFTSGVTNNGRD 171
OmpE36: DTGV-SDNFFSGRTGGLATYRNSGFFGLVDGLNFGVQYLGKNER--------------TD 160
OmpK36: DTYG-SDNFLQSRANGVATYRNSDFFGLVDGLNFALQYQGKNGSVSGEG----ATNNGRG 168

OmpF:   ARRSNGDGVGGSISYE-YEG--FGIVGAYGAADRTNLQEAQ-PLGNGKKAEQWATGLKYD 221
OmpC:   ALRQNGDGVGGSITYD-YEG--FGIGGAISSSKRTDAQNTAAYIGNGDRAETYTGGLKYD 228
OmpE36: ALRSNGDGWATSLSYD-FDG--FGIVGAYGAADRTNAQQNL-QWGKGDKAEQWATGLKYD 216
OmpK36: WSKQNGDGFGTSLTYDIWDG--ISAGFAYSHSKRTDEQNSVPALGRGDNAETYTGGLKYD 226

OmpF:   ANNIYLAANYGETRNATPITNKFTNTSGFANKTQDVLLVAQYQFDFGLRPSIAYTKSKAK 281
OmpC:   ANNIYLAAQYTQTYNATRVG-----SLGWANKAQNFEAVAQYQFDFGLRPSLAYLQSKGK 283
OmpE36: ANNIYLAALYGEMRNAARLD------NGFANKTQDFSVVAQYQFDFGLRPSIAYYKSKAK 270
OmpK36: ANNIYLASQYTQTYNATRAG-----SLGFANKAQNFEVVAQYQFDFGLRPSVAYLQSKGK 281

OmpF:   DVE-GIGDVDLVNYFEVGATYYFNKNMSTYVDYIINQIDSDN---KLGVGSDDTVAVGIV 337
OmpC:   NLGRGYDDEDILKYVDVGATYYFNKNMSTYVDYKINLLDDNQFTRDAGINTDNIVALGLV 343
OmpE36: DVE-GIGDEDYINYIDIGATYYFNKNMSTYVDYQINQLKDDN---KLGINNDDTVAVGLV 326
OmpK36: DLERGYGDQDILKYVDVGATYYFNKNMSTYVDYKINLLDDNSFTRNAGISTDDVVALGLV 341

OmpF:   YQF 340
OmpC:   YQF 346
OmpE36: YQF 329
OmpK36: YQF 344
```

Figure S13: Sequence alignment of OmpF (*E. coli*), OmpC (*E. coli*), OmpK36 (*K. pneumoniae*) and OmpE36 (*E. cloacae*). Each residue is represented by their one-letter code. Important residues in the observed gating mechanism of OmpF are highlighted with a box and colored based on residue type (black: hydrophobic, green: polar, red: acidic, and blue: basic). All other residues are colored gray. The alignment was performed on full-length sequences using the Clustal Omega[87] alignment tool within Uniprot.[88]
